# Curcumin and NCLX Inhibitors Share Anti-Tumoral Mechanisms in Microsatellite-Instability-Driven Colorectal Cancer

**DOI:** 10.1101/2022.01.18.476775

**Authors:** Maxime Guéguinou, Sajida Ibrahim, Jérôme Bourgeais, Alison Robert, Trayambak Pathak, Xuexin Zhang, David Crottès, Jacques Dupuy, David Ternant, Valérie Monbet, Roseline Guibon, Hector Flores-Romero, Antoine Lefèvre, Stéphanie Lerondel, Alain Le Pape, Jean-François Dumas, Philippe G. Frank, Alban Girault, Romain Chautard, Françoise Guéraud, Ana J. García-Sáez, Mehdi Ouaissi, Patrick Emond, Olivier Sire, Olivier Hérault, Gaëlle Fromont-Hankard, Christophe Vandier, David Tougeron, Mohamed Trebak, William Raoul, Thierry Lecomte

## Abstract

Colorectal cancer (CRC) is associated with high mortality worldwide and new targets are needed to overcome treatment resistance. Recent evidences highlight a role of the mitochondria calcium homeostasis in the development of CRC. In this context, we aimed to evaluate the role of the mitochondrial sodium-calcium-lithium exchanger (NCLX) and its targeting in CRC. We also identified curcumin as a new potential inhibitor of NCLX.

In vitro, curcumin exerted strong anti-tumoral activity through its action on NCLX with mtCa2+ and reactive oxygen species overload associated with a mitochondrial membrane depolarization, leading to reduced ATP production and apoptosis through mitochondrial permeability transition pore opening concomitant with G2/M cell cycle arrest. NCLX inhibition with either CGP37157 (a benzodiazepine derivative), small interfering RNA-mediated knock-down or knockout approaches reproduced the effects of curcumin. Altered mitochondrial respiration, cellular aerobic glycolysis and endoplasmic reticulum–mitochondria membrane perturbations participated in these mechanisms. In a xenograft mouse model, NCLX inhibitors decreased CRC tumor growth. Both transcriptomic analysis of The Cancer Genome Atlas dataset and immunohistochemical analysis of tissue microarrays from 381 patients with microsatellite instability (MSI)-driven CRC demonstrated that higher NCLX expression was associated with MSI status and for the first time NCLX expression was significantly associated with recurrence-free survival in MSI CRC patients.

Our findings provide strong evidence that blocking NCLX inhibits CRC in vitro and in vivo. We highlight a novel anti-tumoral mechanism of curcumin through its action on NCLX and mitochondria calcium overload that could benefit for therapeutic treatment of patients with MSI CRC.

## Introduction

Colorectal cancer (CRC) is the third most common cancer worldwide and remains a major public health issue. CRC development is a multi-step process involving distinct stages: initiation, promotion and progression that ultimately generates phenotypically altered transformed malignant cells. Understanding the mechanisms that drive tumor hallmarks is critical for the prevention and treatment of CRC progression using drugs acting on these identified mechanisms. Based on the current state of knowledge, CRC develops mainly through a progressive accumulation of genetic alterations by three major mechanisms. The most common is chromosomal instability (CIN) in 75% of CRCs; the second most common is epigenetic modification of DNA methylation, also called the CpG island methylator phenotype (CIMP), in 20% of CRCs; the third most common is microsatellite instability (MSI) or deficiency of DNA mismatch repair system (dMMR), which occurs in approximately 15% of CRCs (1). MSI CRC cell lines are more resistant to 5-fluorouracil (5-FU) compared with microsatellite stable (MSS) cell lines (2). Indeed, determination of the dMMR/MSI status in CRC has prognostic and therapeutic implications. Additionally, MSI is associated with a better prognosis in early-stage CRC but a worse prognosis at advanced stages (4).

Several preclinical studies support the anti-tumoral properties of curcumin against CRC (5)(6). Curcumin, also known as turmeric or diferuloylmethane, is a natural yellow-pigmented polyphenol extracted from the plant *Curcuma longa*. This compound is a highly pleiotropic molecule with numerous targets and mechanisms of action and modulates several pathways (5). Curcumin is viewed as a potential CRC chemopreventive agent because of its effect on multiple tumor progression pathways (7). Chemoprevention is a promising strategy for reducing CRC incidence. In CRC, the MMR system strongly influences the sensitivity of cells to curcumin. Specifically, the MMR system modulates curcumin sensitivity through induction of DNA strand breaks and activation of the G2/M checkpoint (8). In tumor cells, curcumin participates in mitochondria-dependent apoptotic processes (9). Curcumin affects mitochondria-associated proteins, such as Bax, Bcl-2 and Bcl-xL, and reactive oxygen species (ROS) production (10). Curcumin increases rat liver mitochondrial membrane permeability, resulting in swelling, loss of membrane potential and inhibition of adenosine triphosphate (ATP) synthesis (11). Recently, intracellular distribution of curcumin has been correlated with mitochondrial staining (12). Notably, Jelinek and colleagues demonstrated a preferred association of curcumin with cardiolipins, lipids mainly localised in the inner mitochondrial membrane (12). Despite an abundant literature concerning curcumin and its effects, the molecular mechanisms of action remain poorly described, in particular the link between mitochondrial redox status and mitochondria-associated redox and calcium (Ca^2+^) signaling mechanisms.

The interplay between Ca^2+^ and ROS signaling pathways is well established at several subcellular locations and has been implicated in many essential biological functions (13)(14). Mitochondrial permeability transition pore (mPTP) opening is triggered by different pathological conditions such as Ca^2+^ overload and oxidative stress (15). This opening leads to reduced ATP production and allows the release of pro-apoptotic mitochondrial components, namely apoptosis-inducing factor and cytochrome *c*. Another hallmark of mPTP-induced apoptosis is the loss of mitochondrial membrane potential. Several proteins participate in mitochondrial Ca^2+^ (mtCa^2+^) uptake:, the mitochondrial calcium uniporter (MCU) and its known regulators, which currently include MICU1, MICU2, EMRE, MCUR1 and MCUb. The mitochondrial Na^+^/Ca^2+^/Li^+^ exchanger (NCLX) mostly mediates mtCa^2+^ extrusion (16). Of note, NCLX has been underexplored as a potential therapeutic target to restore abnormal Ca^2+^ fluxes in cancer (17)(18). In CRC, we have recently shown that downregulation of NCLX results in mtCa^2+^ overload, increased mitochondrial ROS (mtROS), mitochondrial depolarisation and tumor shrinkage (19). The benzodiazepine-derived compound CGP37157, an inhibitor of NCLX, promotes mitochondrial damage and induces apoptosis (20).

In the present study, we examined whether curcumin induces a modification of mtCa^2+^ flux in association with redox metabolism in MSI CRC. Our results indicated that curcumin has a strong anti-tumoral effect through inhibiting NCLX: inhibition of G2/M cell cycle progression, mtCa^2+^ overload, mitochondrial membrane depolarization, production of ROS leading to reduced ATP production and apoptosis by mPTP opening in MSI CRC cells. In a xenograft mouse model of CRC, NCLX inhibitors decreased CRC growth. In addition, mid-infrared (MIR) spectroscopy coupled with metabolomics gave useful metabolic signatures in terms of circulating biomarkers. Moreover, analysis of CRC in The Cancer Genome Atlas (TCGA) dataset showed that higher *NCLX* expression was associated with MSI status, and the expression of NCLX in tissue microarrays (TMA) of 302 samples was significantly associated with recurrence-free survival (RFS) in patients with MSI CRC.

## Results

### Curcumin Inhibited Proliferation and Induced Cell Cycle Arrest of Colorectal Cancer Cells

We investigated the role of curcumin in regulating the biology of CRC cells. Curcumin (0–10 µM) dose-dependently reduced the viability of these cells based on the 3-(4,5-dimethylthiazol-2-yl)-2,5-diphenyl-2H-tetrazolium bromide (MTT) assay (Figure 1A) and the Sulforhodamine B (SRB) assay (Supplementary Figure 1A). Moreover, 10µM curcumin decreased the proliferation of HCT116 cells, a CRC cell line, for up to 72 h (Figure 1A, left). This pattern was identical in two other MSI cell lines, Lovo and SW48 cells (Supplementary Figure 1B). Interestingly, curcumin had no effect on the proliferation of a non-tumorigenic epithelial colorectal cell line, NCM356 (Figure 1A right). We next examined whether curcumin affected the proliferation of CRC cells by using the spheroid three-dimensional (3D) assay. There was a significant reduction in HCT116 and Lovo multicellular spheroid volume with respect to the control group after administration of 10 µM curcumin for 48 h (Figure 1B) confirmed by the measure of the cell viability index (Figure 1C). Cell-cycle analysis revealed that curcumin induced a large, dose-dependent increase in the percentage of HCT116 and SW48 cells in the G2/M phase (Figure 1D) suggesting that curcumin can reduce cell proliferation by preventing mitosis.

**Figure 1:**
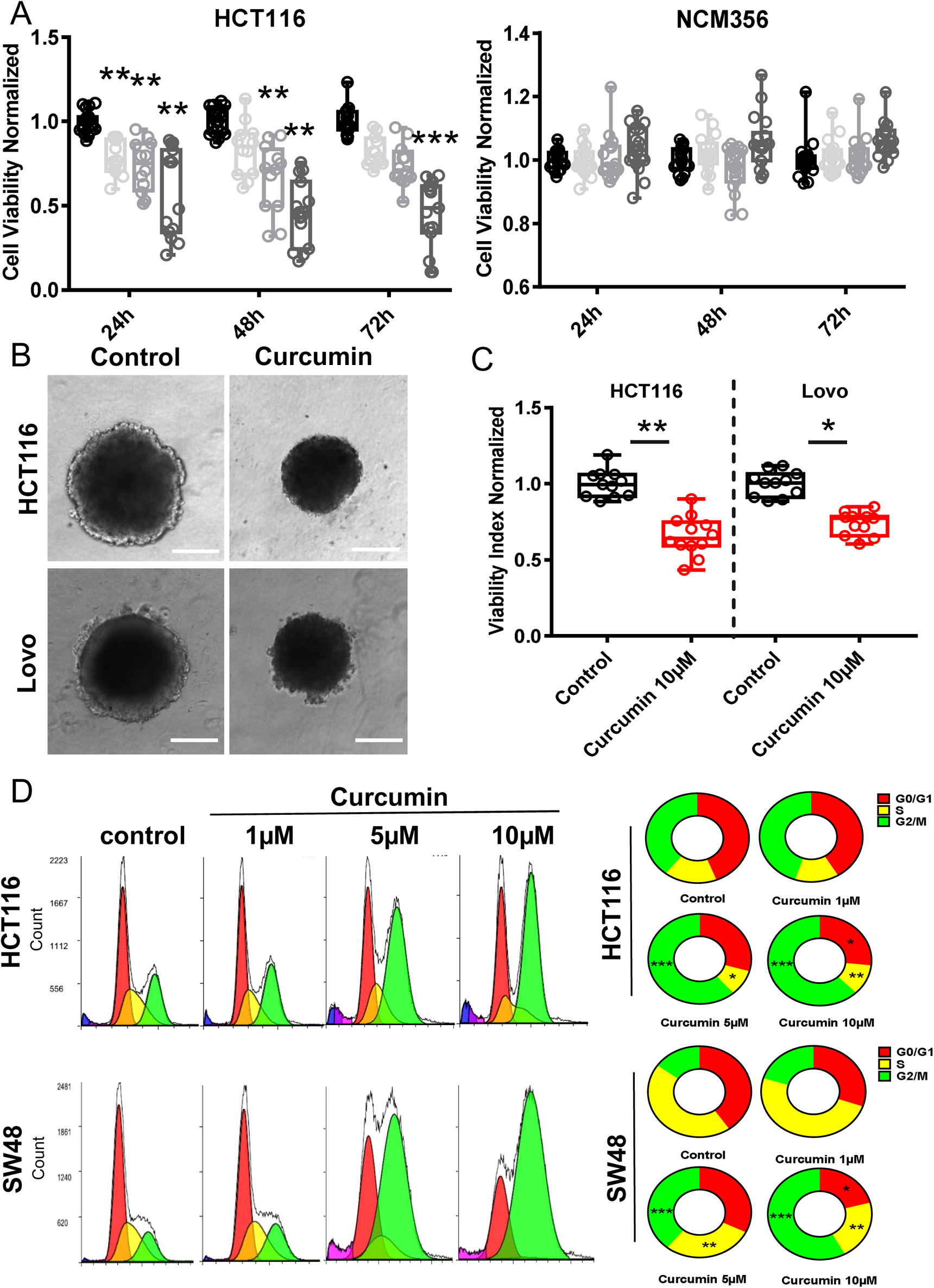
Curcumin exerts anti-proliferative effects on human CRC cells by inducing G2/M cell cycle arrest. A/ Dose-dependent effect of curcumin on cell viability of normal human colorectal cells and human CRC cells. Human colorectal cells (NCM356) and human MSI CRC cells (HCT116) were incubated with various concentrations of curcumin (0–10 µM) for 24, 48 or 72 h, and cell viability was measured using the MTT or SRB assay (supplementary data Figure 1). The results are expressed as the mean ± standard deviation of 3–5 independent experiments (ANOVA followed by Tukey’s post hoc multiple comparisons, \*\**P* < 0.01 and *** *P* < 0.001,). B/ The effect of curcumin on 3D spheroids. Cell proliferation with the Cultrex 3D proliferation assay of Lovo and HCT116 spheroids exposed to 10 µM curcumin for 72 h. The area of 3D spheroids after incubation with 10 µM curcumin (72 h) was measured using the ImageJ software. C/ The effect of curcumin on the cell viability index of spheroids The cell viability index was evaluated by measuring resofulvin reduction in HCT116 and Lovo cells (n = 5 independent experiments; Mann–Whitney test, * *P* < 0.05, ** *P* < 0.01). D/ Cell cycle evaluation. Curcumin triggers G2/M cell cycle arrest in CRC cells as measured by flow cytometry using the propidium iodide assay (n = 5 independent experiments; ANOVA followed by Tukey’ post hoc multiple comparisons, \*\**P* < 0.01 and \*\*\**P* < 0.001).

### Curcumin Inhibited Mitochondrial Ca^2+^ Extrusion and Induced Reactive Oxygen Species Production with Mitochondrial Membrane Potential Depolarization

Cell proliferation is controlled by intracellular Ca^2+^ and ROS, with mitochondria acting as a Ca^2+^ buffer and ROS generators (14). We measured mtCa^2+^ in HCT116 cells by loading the cells with Rhod-2AM, a mtCa^2+^-sensitive dye. Curcumin treatment for 1 hour induced a sharp increase in the mtCa^2+^ concentration under basal conditions (Figure 2A). Cells were then stimulated with ATP, a purinergic G protein-coupled receptor (P2Y) agonist that is coupled to phospholipase Cβ activation and subsequent inositol-1,4,5-trisphosphate (IP3) production. IP3 binds to IP3 receptors (IP3R) and then mediates Ca^2+^ transfer from the endoplasmic reticulum (ER) to the mitochondria. Curcumin had no effect on mtCa^2+^ uptake but inhibited mtCa^2+^ efflux after stimulation with 100 µM ATP (Figure 2B). Recently, we have shown that NCLX is the most important actor of mtCa^2+^ extrusion in CRC cells and that inhibiting NCLX expression and function prevents mtCa^2+^ extrusion without affecting mtCa^2+^ uptake (Supplementary Figure 2A – 2B)(19). To confirm these effects on NCLX we performed mtCa^2+^ uptake and release experiments Na^+^ dependent on permeabilized HCT116 cells (Supplementary Figure 2C-D). Permeabilized HCT116 cells suspended in intracellular Na^+^-free buffer. As expected, a slow Ca^2+^ extrusion was observed after Na^+^ injection in CGP37157 and curcumin treated cells without additive effect. The knockdown of either NCLX function (through the over-expression of the S486T mutant, described as a dominant-negative) or expression significantly reduced the effect of curcumin on cell viability (Figure 2C). Interestingly, the pharmacological inhibition of NCLX, using CGP37157, in addition to curcumin during 24h did not reduce the cell viability when compared to either curcumin or CGP37157 alone suggesting that both compounds are acting on the same mechanisms or the same target (in Figure 2D). Altogether, these suggest that curcumin effect on cell viability are mediated by an effect on the NCLX-dependent mtCa^2+^ efflux, Through the modulation of Na^+^ influx and Ca^2+^ efflux, NCLX directly acts on the mitochondrial potential (19). Here, we observed that both curcumin and CGP37157 induce a depolarization in HCT116 as visualized through the measure of the fluorescence of the mitochondrial potential reporter dye tetramethylrhodamine methyl ester (TMRE). Interestingly, the addition of curcumin and CGP37157 during 1 hour does not produce additional depolarization arguing that both drugs modulate mitochondrial potential through NCLX (Figure 2E with TMRE).

**Figure 2:**
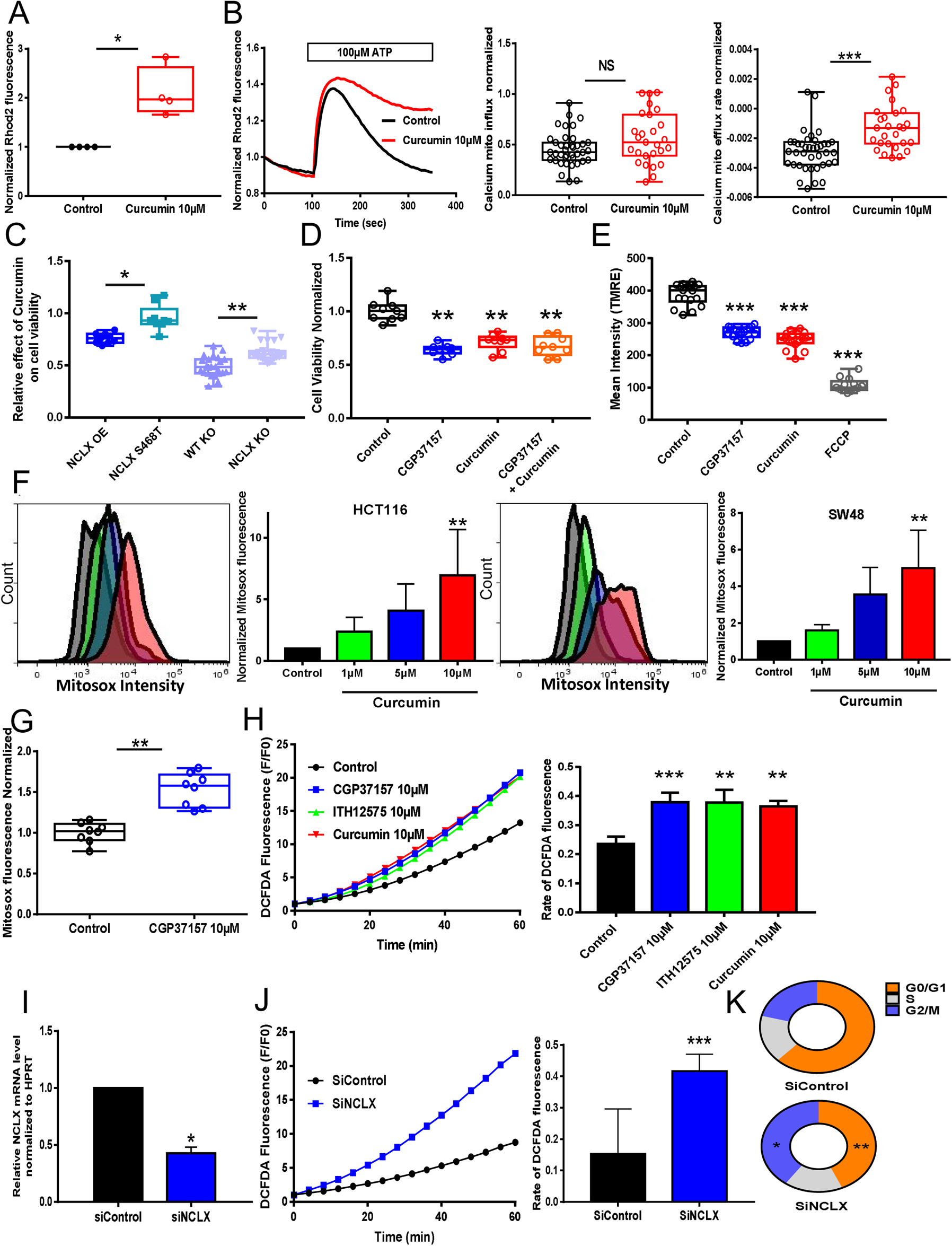
mtCa^2+^ efflux and ROS production are altered by treatment with curcumin and NCLX targeting in CRC cells. A/ Global mtCa^2+^ was evaluated with Rho2 fluorescence in HCT116 cells treated or not treated with curcumin (n = 4 independent experiments; Mann–Whitney test, * *P* < 0.05). B/ mtCa^2+^ responses following ATP stimulation (100 μM) were measured in HCT116 cells treated or not treated with curcumin. The average mtCa^2+^ influx (peak) (left) and the rates of mtCa^2+^ efflux (right) are shown (n = 5 independent experiments with 27–39 measurements; Mann–Whitney test, *** *P* < 0.001). C/ The knockdown of either NCLX function (through the over-expression of the S486T mutant, described a dominant-negative) or expression significantly reduced the effect of curcumin on cell viability. The effects of curcumin were reduced in HCT116 (n=4 independent experiments of 9 measurements per Group over express NCLX or NCLX S468T \**P* < 0.05,n = 5 independent experiments of 24 measurements per group for WT KO or NCLX KO cells, \*\**P* < 0.01) C/ mtCa^2+^ responses following application of ATP (100 μM, as in Figure 3A) were measured in HCT116 control (NCLX +/+) or NCLX KO (−/−) cells. The average mtCa^2+^ influx (peak) (left) and rates of mtCa^2+^ efflux (right) are shown (n = 9, Mann-Whitney test, \*\**P* < 0.01) D/ CGP37157 and curcumin had no additive effect on cell viability (MTT assay). The results are expressed as the mean ± standard deviation of 3–5 independent experiments (ANOVA followed by Tukey’s post hoc multiple comparisons, \**P* < 0.05, \*\**P* < 0.01). E/ Mitochondrial membrane potential using the dye TMRE in HCT116 cells treated with M or CGP37157 10 μM OR FCCP 50µM. (n =6, Mann-Whitney test, \*\*\**P* < 0.01). F/ HCT116 and SW48 cells were pre-treated with curcumin (1, 5 or 10 μM) for 2 h. The cells were stained with MitoSOX for 40 min. The bar graph presents a quantitative analysis of mtROS. The data represent the mean ± standard deviation of four independent experiments (n = 4 independent experiments, \*\**P* < 0.01, ANOVA followed by Tukey’s post hoc multiple comparisons). G/ The effect of CGP37157 pre-treatment on the mtROS level in HCT116 cells was recorded. The data are shown as the mean ± standard error of the mean of at least three separate experiment with eight measurements each (n=8, Mann-Whitney test, \*\**P* < 0.01). H/ Cytosolic ROS is indicated by an increase in DFDCA fluorescence over time. DCFDA fluorescent traces are shown as the normalised (F/F0) mean. The effects of curcumin and NCLX blockers (CGP37157 or ITH12575) on cytosolic ROS responses in HCT116 cells were analysed. The histograms show the slope of cytosolic ROS production as a function of time (n = 5 independent experiments with 31–56 measurements per group \*\**P* < 0.01, \*\*\**P* < 0.001; ANOVA followed by Tukey’s post hoc multiple comparisons). I/ RT-qPCR data showing *SLC8B1* mRNA relative to *HPRT* in HCT116 cells transfected with siRNA against NCLX (n = 4 independent experiments, Mann-Whitney test, \**p* < 0.05) J/ The effect of siRNA-mediated NCLX inhibition on cytosolic ROS production was evaluated in HCT116 cells (n = 4 independent experiments with 16–32 measurements per group, Mann-Whiney test, \*\**P* < 0.01). K/ Cell cycle analysis by flow cytometry. Cells were transfected with siRNAs. After 48 h, HCT116 cells were fixed and stained with propidium iodide. The results are representative of five independent experiments.

A further consequence of mtCa^2+^ overload is the generation of mtROS. Using mitoSOX dye to monitor the generation of mtROS, we observed that curcumin induces a dose-dependent increase of mtROS in HCT-116 and SW48 cell lines. (Figure 2F and Supplementary 2C). 5-FU treatment and pharmacological inhibition of NCLX with CGP37157 also increased mtROS in a same range as curcumin (Figure 2F and supplementary 2E).

All treatments for 1h including curcumin, pharmacological inhibitors of NCLX (CGP37157 or ITH12575) and downregulation of NCLX with siRNA not only increased mtROS but also enhanced cytosolic ROS demonstrated by the measure of DCFDA (Figure 2G–J). Here, we observed that the silencing of NCLX expression can also impair cell cycle similarly to curcumin (Figure 2K). Overall, our results indicate that curcumin may regulate mtCa^2+^ efflux and ROS production through the inhibition of NCLX.

### Curcumin and NCLX Inhibition Decreased Store-Operated Calcium Entry

In non-excitable cells, the intracellular Ca^2+^ concentration is controlled by store-operated calcium entry (SOCE) through CRAC, which are encoded by Orai proteins located at the plasma membrane of CRC cells. This influx is important for several cellular processes, including proliferation, migration and invasion (21). NCLX inhibition through siRNA knockdown in CRC cells and in HEK293 cells leads to the inhibition of plasma membrane Orai1 channels and reduces SOCE (19)(22). Here, using the whole cell patch-clamp technique, we observed that curcumin abolished the Orai1-mediated I_CRAC_ currents induced by Thapsigargin (Tg) in an heterologous expression system co-expressing STIM1 and Orai-1 (Figure 3A). In addition, curcumin pre-treatment for 1 hour reduced SOCE in HCT116 cells (Figure 3B). To identify the mechanism involved in regulation of SOCE, the messenger RNA (mRNA) levels of the Ca^2+^ handling proteins *MCU*, *NCLX*, *MICU1*, *MICU2*, *STIM1*, *STIM2*, *Orai1* and *TRPC1,* only *MICU1* was significantly increased after 24h treatment with curcumin suggesting that its effect is not mediated by transcriptional mechanisms (Figure 3C). To determine whether curcumin-dependent changes in SOCE and CRAC are mediated by a change in STIM1 oligomerisation, we monitored the formation of STIM1-RFP puncta in HCT116 cells after Tg-induced ER Ca^2+^ depletion. Indeed, oligomerisation of STIM1 coupled with ER Ca^2+^ depletion is necessary for CRAC activation. Numerous puncta were formed following Tg stimulation and curcumin had no significant effect (Figure 3D). Interestingly, this reduced Ca^2+^ influx induced by curcumin was also observed in NCLX KO cells (Figure 3E) or after CGP37157 treatment (Figure 3F). As expected, molecular silencing and pharmacological inhibition of NCLX reduced Ca^2+^ influx in a similar manner than curcumin.

**Figure 3:**
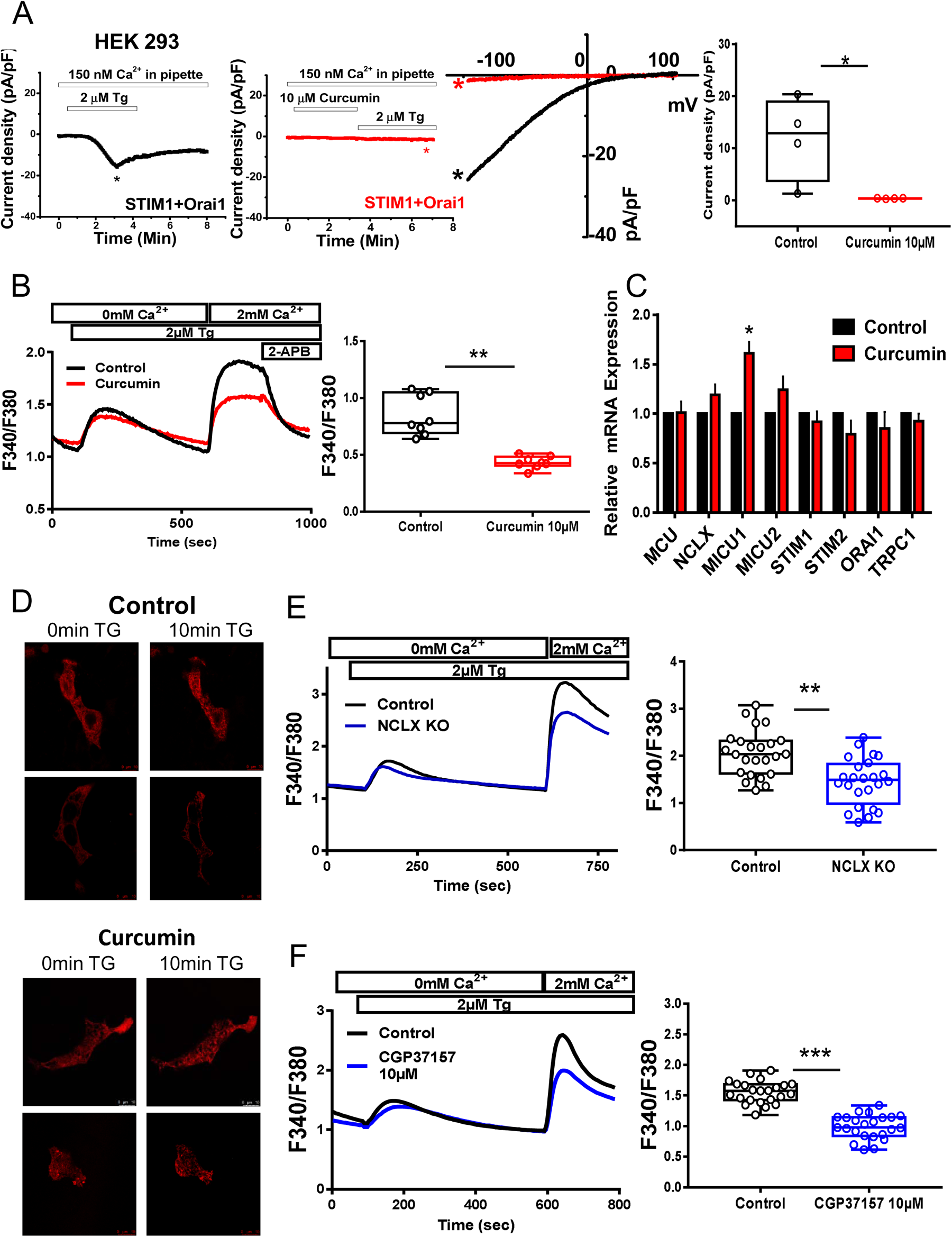
The effect of NCLX inhibition and curcumin treatment on SOCE in CRC cells. A/ Electrophysiological recordings were performed on HEK293 ORAI1 KO cells transfected with ORAI1-CFP and STIM1-YFP. Representative time courses of whole-cell CRAC currents activated by dialysis of 20 mM BAPTA through the patch pipette and taken at −100 mV from control cells (black trace) or cells pre-treated with 10 µM curcumin (red trace) are shown. Representative I–V relationships are taken from traces in (Figure 3A) are indicated by colour coded asterisks. Statistical analysis of Ca^2+^ CRAC currents measured at −100 mV is shown (n = 4 independent experiments; Mann-Whiney test, *P < 0.05). B/ Representative traces of SOCE in HCT116 cells pre-treated with curcumin (10 µM for 1 h). The data show the effects of curcumin on SOCE in HCT116 (n = 5 independent experiments; Mann-Whiney test, \*\**P* < 0.01). C/ The relative mRNA levels of *MCU*, *NCLX*, *MICU1*, *MICU2*, *STIM1*, *STIM2*, *ORAI1* and *TRPC1* in HCT116 cells treated or not treated with 10 µM curcumin for 24 h (n = 5 independent experiments; Mann-Whiney test, \**P* < 0.05). D/ STIM1-dependent puncta formation a. Representative fluorescent images of HCT116 cells co-expressing STIM1 Cherry (red) showing puncta after treatment with TG (2 µM for 10 min) and pre-treated or not pre-treated with curcumin for 30 min. E/ Representative cytosolic Ca^2+^ measurements in HCT116 WT and HCT116 NCLX KO cells measured with Fura-2 AM in response to 2 µM TG applied first in nominally Ca^2+^-free external solution and subsequently with 2 mM external Ca^2+^. The summary data are presented as the mean ± standard error of five independent experiments with 24 cells for each group (n = 5 independent experiments; Mann–Whitney test, \*\**P* < 0.01). F/ SOCE was blocked after preincubation with CGP37157 (10 μ for 1 h) (right graph). Ca influx in control and CGP37157-treated cells is shown. The summary data are presented as the mean ± standard error of the mean of five independent experiments with 23 cells for each group (n = 4 independent experiments; Mann–Whitney test, \*\*\**P* < 0.001).

### Curcumin and NCLX Inhibition Altered Colorectal Cancer Metabolism

Cancer cell proliferation and mitosis are highly dependent of the cellular energy produced by mitochondria through mitochondrial oxidative phosphorylation (23). mtCa^2+^ and mtROS are key players in the control of mitochondrial metabolism and ATP production. Because Ca^2+^ regulates key metabolic enzymes, we hypothesised that by targeting NCLX, curcumin or CGP37157 affects mtROS and mtCa^2+^ to limit cellular bioenergetics. To test this hypothesis, we determined the effects of these compounds on mitochondrial respiration (OXPHOS) and aerobic glycolysis (Figure 4A–C). Both curcumin and CGP37157 treatment reduced the intact cellular oxygen consumption rate (OCR) in HCT116 cells; this change was associated with a reduction in basal respiration, maximal respiration, proton leak and ATP production (Figure 4C, left). By contrast, curcumin and CGP37157 had no significant effects on glycolysis, glycolytic capacity and glycolytic reserve (Figure 4C, right). Hypoxia significantly increased the expression of HIF1α compared with normoxia. Under both conditions, NCLX inhibition by curcumin or CGP37157 significantly reduced the activity of HIF1α (Figure 4D). Surprisingly, we found an opposite effect in cells transfected with siRNA against NCLX. In this condition, the reduction in NCLX increased HIF1α in both normoxic and hypoxic conditions (Figure 4D). Pathak et al. (19) showed that inhibition of NCLX expression with either short hairpin (shRNA) or CRISPR/Cas9 knockout enhances HIF1a activation and increases the ECAR of HCT116 cells with an increase in glycolysis. Altogether, these data suggest that curcumin may affect cancer cell viability by its action on NCLX and impairing mitochondrial respiration.

**Figure 4:**
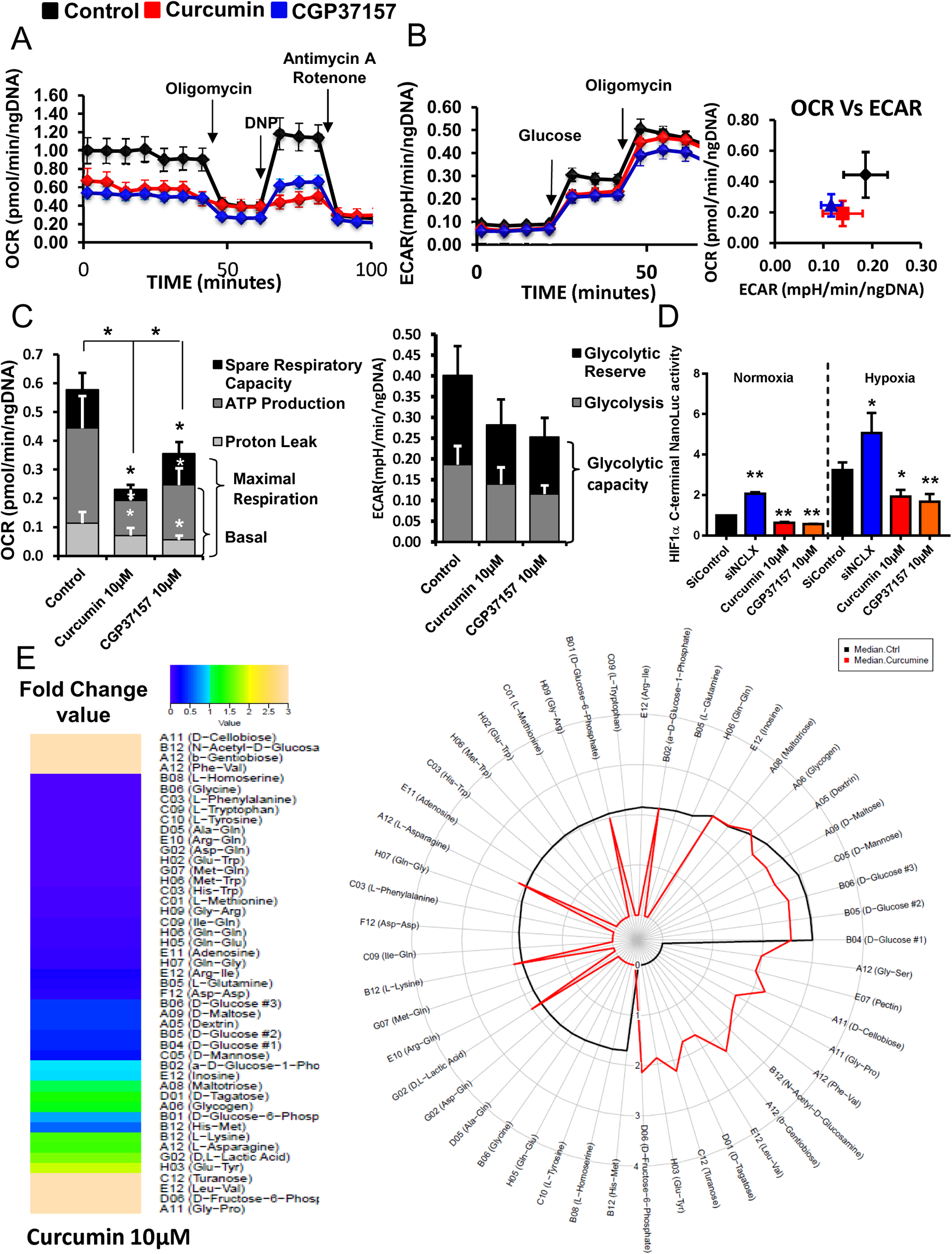
Targeting NCLX-dependant signaling elicits rapid and dynamic changes in CRC cell metabolism. A/ Energy metabolism in HCT116 cells pre-treated with curcumin and NCLX blocker (CGP37157). After four measurements of baseline OCR, glucose, oligomycin, DNP and the mixture of antimycin A and rotenone were injected sequentially, with measurements of OCR recorded after each injection. ATP-linked OCR and OCR due to proton leak can be calculated using basal and oligomycin-sensitive rates. Injection of DNP is used to determine the maximal respiratory capacity. Injection of antimycin A and rotenone allows for the measurement of OCR independent of complex IV. B/ Cellular aerobic glycolysis was evaluated by measures of the ECAR from HCT116 cells treated with curcumin or CGP37157 (10 µM) following sequential injection (arrow) of glucose (10 mM) and then oligomycin (1 μmol/L) (n = 5 independent experiments; Mann–Whitney test, \*\*\**P* < 0.001). C/ The spare respiratory capacity (mitochondrial reserve capacity) was calculated by subtracting the basal from the maximal respiration. The bioenergetic profiles of HCT116 with or without 10 µM curcumin or 10 µM CGP37157 were measured using sequential injection of glucose, oligomycin (1 μmol/L), DNP (100 μmol/L) and the mixture of antimycin A (0.5 μmol/L). The mitochondrial reserve capacity of untreated control and curcumin- or CGP37157-treated HCT116 cells are shown. The results are presented as the mean ± standard deviation of five independent experiments of 3–5 measurements Mann–Whitney test, \**P* < 0.05, \*\*\**P* < 0.001). D/ HCT116 HIF1α-C-terminal Luc cells incubated under normoxic or hypoxic conditions for 48 h. Four hours before the end of incubation, the cells were treated with CGP37157 or curcumin. The values represent the mean ± standard deviation of 3–6 independent experiments. E/ The effect of curcumin on the ability of HCT116 cells to metabolise 367 substrates was measured by using the OmniLog® Analyzer. The heatmap shows the AUC fold-change data induced by curcumin treatment. The radial plot shows the AUC of metabolised substrates (49 substrates with positives kinetic curves with AUC > 50 in at least one conditions) on a logarithmic scale.

Previous transcriptomic analysis showed that glycolysis-related genes are upregulated in HCT116 NCLX KO cells (19). To measure the impact of curcumin on the remodeling of the metabolism, we performed real-time quantitative PCR on a set of 76 genes associated with metabolism. Confirming the bioenergetic results, curcumin treatment had a moderate effect on glycolysis-related genes or antioxidant genes (Supplementary Figure 4). In terms of energy metabolism, there was a significant decrease in the expression of Glut1 (SLC2A1) and Glut4 (SLC2A4) transporters, as well as hexokinase 2, associated with an increase in glucokinase regulator (GCKR) expression and a decrease in PFKFB1 and B4 (glycolysis activator) expression. These changes favour an overall decrease in carbohydrate metabolism. We also observed a decrease in MCT2 (lactate import) and lactate dehydrogenase A transcripts. Transcript quantification showed an increase in MCT4 (lactate export) and lactate dehydrogenase C associated with the decrease in PDHB (pyruvate dehydrogenase E1 complex or PDC-E1) and a drop in PDP1 (PDC activator) as well as PDC inhibitors (Supplementary Figure 4). Taken together, these results strongly suggest that curcumin decreases the use of pyruvate for oxidative phosphorylation and increases the lactate efflux. However, in the context of complexes such as PDC, we cannot make assumptions about enzymatic activities. With regard to oxidative metabolism, curcumin induced a decrease in GLRX5 and PRDX3 expression and an increase in the GPX family of enzymes (Supplementary Figure 4). However, these genes were poorly expressed, suggesting that they would have a moderate effect on ROS levels. Finally, curcumin increased the expression of nuclear factor (erythroid-derived 2)-like 2 (NRF2) target genes (Supplementary Figure 4). This phenomenon could explain the presence of oxidative stress. NRF2 expression was enhanced following curcumin treatment in normoxic conditions and following curcumin or siNCLX treatment in hypoxic conditions (Supplementary Figure 3A). However, these treatments did not affect the activity of NRF2 induced by its nuclear translocation (Supplementary Figure 3B).

To better characterise the metabolic effects of curcumin, we used the OmniLog® Phenotype MicroArray, that allows to generate a map of the metabolism phenotype of cancer cells. Treating HCT116 cells with curcumin blocked the metabolism of L-glutamine, L-tryptophan, L-methionine, adenosine, L-phenylalanine, glycine, L-tyrosine, L-homoserine and different dipeptides (Figure 4E). By contrast, b-gentiobiose, D-cellobiose, turanose, D-tagatose, D-fructose-6-phosphate, *N*-acetyl-D-glucosamine pectin and different dipeptides were only used by HCT116 cells treated with curcumin (Figure 4E). In addition, 14 substrates were found in both curcumin-treated and non-treated HCT116 cells. Finally, curcumin induced a decrease in the use of major substrates, both in terms of biological importance and level of use, such as D-glucose (−60%), D-mannose (−69%), D-maltose (−58%), dextrin (−55%) and D-glucose-6-phosphate (−20%), whereas there was an increase in the consumption of substrates such as glycogen (+21%), L-asparagine (+46%), L-lysine (+48%) and D- and L-lactic acid (+68%).

Taken together, these results suggest that NCLX inhibition with curcumin suppressed mitochondrial respiration, amino acid metabolism and HIF1α-dependent reduction in glycolytic substrate consumption.

### NCLX Inhibitors Induced Apoptosis via Mitochondrial Permeability Transition Pore Sensitisation and Modification of Endoplasmic Reticulum–Mitochondria Contact Sites

High ROS and Ca^2+^ levels in mitochondria with mitochondria membrane depolarization can trigger cell death following the opening of mPTP (14). Apoptosis evaluated by flow cytometry indicated that curcumin dose-dependently increased apoptosis in HCT116 and SW48 cells (Figure 5A and B). To determine whether NCLX plays a role in Ca^2+^-induced mPTP opening, HCT116 cells were loaded with calcein-AM and cobalt was added to quench the cytosolic fluorescence of calcein. HCT116 cells were pre-treated for 1 hour with either 10 µM of curcumin or 1 µM of cyclosporine A (CsA), an mPTP blocker, and exposed to ionomycin to increase mtCa^2+^ (Figure 5C). Remarkably, ionomycin induced more notable calcein fluorescence quenching in HCT116 cells pre-treated with curcumin or CGP37157 for 1 hour compared to control (DMSO) (Figure 5C, Figure 5E). In addition, ionomycin induced greater quenching of calcein fluorescence in HCT116 NCLX KO cells compared with HCT116 WT cells (Figure 5D). These effects were fully reversed by pre-treatment with CsA, indicating that they are mediated by mPTP (Figure 5E). By using Calcium Green-5N, we showed there was greater mPTP activation in siNCLX-transfected HCT116 cells (Supplementary Figure 3C). Taken together, either NCLX inhibition or curcumin can sensitise CRC cells to mPTP opening and consequently promote cancer cell death.

**Figure 5:**
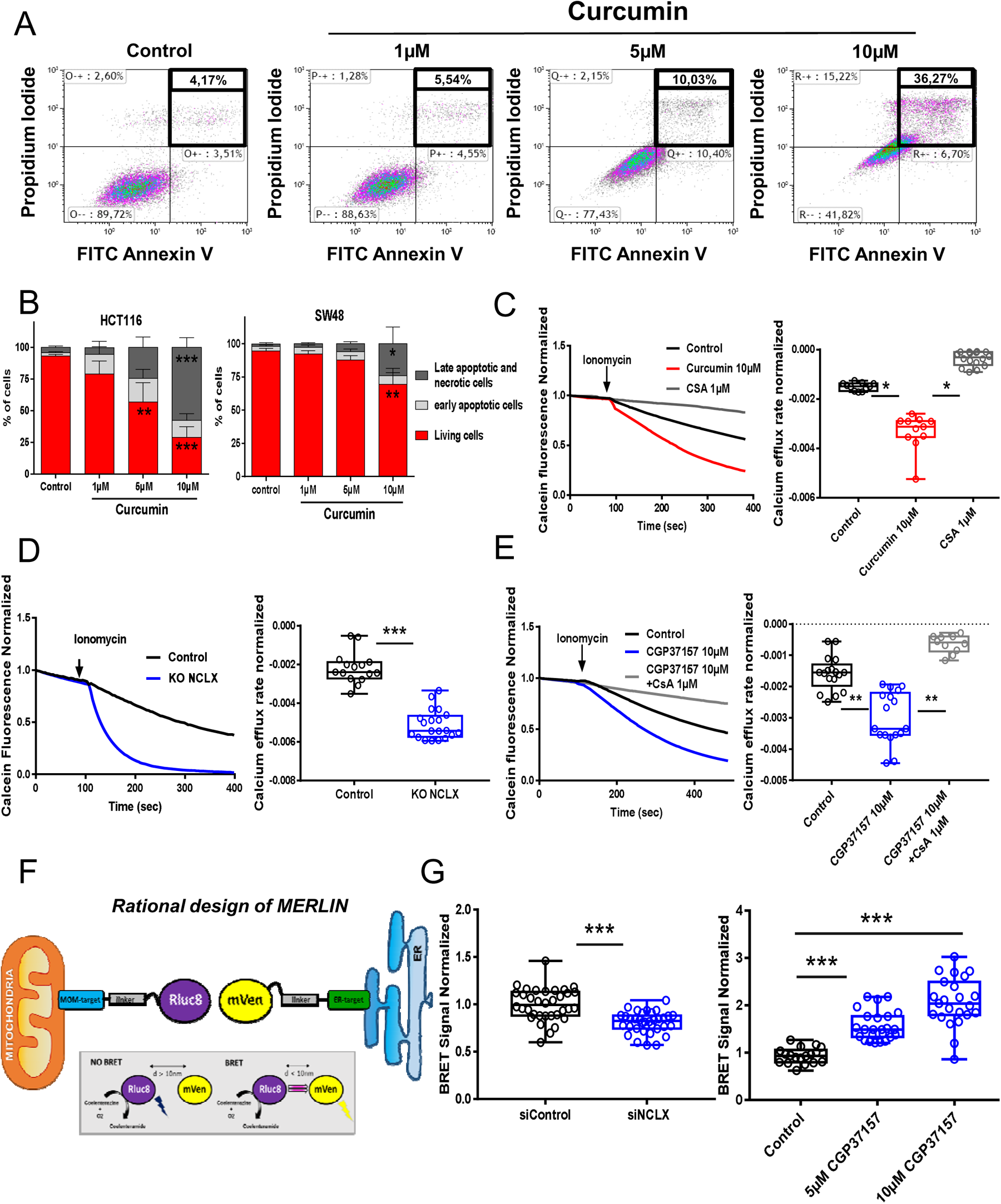
Treatment with curcumin or NCLX inhibitors induced Ca^2+^-dependent apoptosis by sensitising mPTP activation. A/ HCT116 and SW48 cells were treated with 0–10 µM curcumin for 24 h. Apoptotic cells were counted by using the Annexin V/PI assay. The dots indicate the number of Annexin V-/PI- cells (bottom left field, indicates live cells), Annexin V+/PI- cells (bottom right field, indicates early apoptotic cells), Annexin V+/PI+ cells (top right field, indicates late apoptotic cells) and Annexin V−/PI+ cells (top left field, indicates necrotic cells). The percentages of early and late apoptotic cells were quantified. The early-stage apoptotic cells are shown in grey and the late-stage apoptotic and necrotic cells are shown in black. B/ The results are presented as the meanD±Dstandard error of 4–5 independent experiments (\**P*D<D0.05, \*\**P*D<D0.01 and \*\*\**P*D<D0.005 compared with their respective control; #*P*D<D0.05, ##*P*D<D0.01 and ns non-significant). C/ For the mPTP opening experiment, HCT116 cells were loaded with calcein-AM and CoCl_2_. mPTP opening was monitored by the decrease in calcein fluorescence induced by 5 µM ionomycin in HCT116 cells pre-treated for 1 h with DMSO (black line), 10 µM curcumin (red line) or 1 μM CsA (grey line). mPTP opening occurs after the addition of 5 μM ionomycin. D/ Depletion of NCLX expression increased mPTP opening activation in HCT116 cells. Measurement of calcein fluorescence in HCT116 WT and NCLX KO cells after activation of mPTP opening induced by 5 μM ionomycin. The kinetics of mPTP opening was increased when the expression of NCLX was downregulated (Mann-Whitney test, \**P* < 0.05). E/ The CsA pre-treatment prevented the mPTP sensitisation induced by CGP37157 treatment. Changes in mitochondrial calcein fluorescence intensities induced by 5 µM ionomycin were measured in HCT116 cells pre-treated for 1 h with 10 µM CGP37157 or =10 µM CGP37157 and 1 µM CsA (One-way ANOVA with Dunnett’s multiple comparisons test, \**P* < 0.05, \*\**P* < 0.01 and \*\*\**P* < 0.001 versus DMSO). F/ Schematic representation of MERLIN based on BRET. The BRET donor was MOM-targeted Rluc8 and the BRET acceptor was mVenus in the ER. In both BRET partners, the linker region is an alpha helix formed by the A(EAAAK)_4_A motif. BRET occurs upon MOM-Rluc8 stimulation by coelenterazine h and RET to ER-mVenus. G/ Targeting NCLX modified the contacts between the ER and mitochondria. Changes in BRET signals in HCT116 cells co-expressing the BRET pair MERLIN, sCal-L1-RLuc and mVen-L1-B33C after knockdown of NCLX, siRNA control transfection or treatment with CGP37157. The results are based on 3–4 replicates with 18–22 measurements per group. The error bars indicate the standard deviation (\*\*\**P* < 0.001, Mann–Whitney test or One way ANOVA with Dunnett’s multiple comparisons test).

It has increasingly been recognised that the ER and mitochondria communicate at specific membrane contact sites called mitochondria-associated ER membranes (MAM) (24). Perturbation of MAM function and/or structure elevates oxidative stress, irreparably damages mitochondria and reduces mitochondrial respiration. These effects have an impact on mitochondrial ATP production and apoptosis. Thus, we investigated the role of NCLX on ER–mitochondria contact sites using MERLIN, a newly developed BRET-based proximity biosensor, expressed in HCT116 cells (Figure 5F)(25). The MERLIN biosensor allows the detection of changes in distance in the nanometre range between the ER and mitochondria, by measuring the energy transfer between the mitochondrial outer membrane (MOM)-targeted Rluc8 (BRET donor) and the ER-targeted mVenus (BRET acceptor). Silencing NCLX substantially decreased the BRET signal (Figure 5G), indicating a decrease in contact points between the ER and mitochondria (Figure 5G, left). However, pharmacological inhibition of NCLX after 3h pre-treatment strongly increased the BRET signal in a dose-dependent manner, suggesting more intimate contact between the ER and mitochondria under these conditions (Figure 5G, right). This apparent discrepancy could be explained by temporal differences and/or modes of action between the siRNA approach and CGP37157 treatment. Increased interaction between the mitochondrial network and ER during early ER stress conditions has already been described (26) Taken together, these data suggest that inhibition of NCLX can contribute to apoptosis with complete sensitisation of mPTP and to modified architecture and spatiotemporal regulation of ER–mitochondria contact sites.

### NCLX Inhibitors Decreased Colorectal Cancer Growth *In Vivo*

The effects of curcumin and CGP37157 were tested in a CRC-xenografted Swiss nude mouse model (schematic diagram, Figure 6A). As shown in Figure 6B (left panel), CRC xenografts that were treated with vehicle (DMSO, Control Group), curcumin or CGP37157 showed decreased primary tumor growth (tumoral volume) compared with control mice. Based on mathematical modelling of tumor growth (Figure 6B, right), the estimated mean (inter-subject standard deviation) model parameters were k_growth_ = 0.46 day^−1^ (–), Vmax = 336 mm^3^ (0.69) and γ = 0.34. More precisely, tumor volume doubled in 1.5 days and reached a maximum average value of 336 mm^3^ in the control group. Vmax was significantly decreased by 2-fold in the presence of either CGP37157 (*P* = 0.033) or curcumin (*P* = 0.040). In the presence of ITH12575, there was a non-significant trend for decreased Vmax (*P* = 0.081).

**Figure 6:**
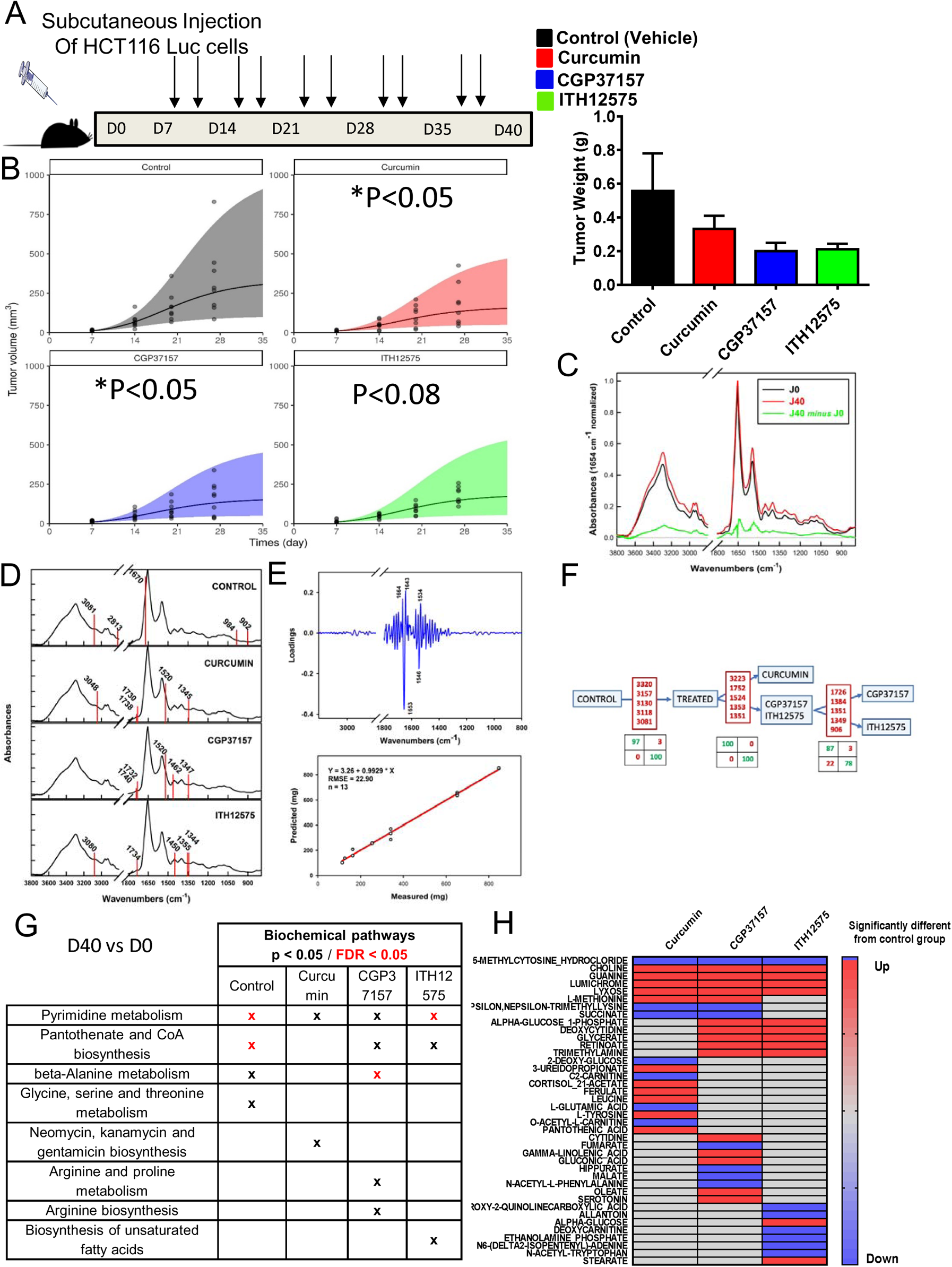
Curcumin and NCLX blockers reduced colorectal tumor growth *in vivo*. A/ HCT116-Luc cells were implanted subcutaneously into nude mice. The mice were divided into four groups: control, curcumin treated, CGP37157 treated and ITH12575 treated (n = 8–9/group). The xenograft volume (tumor volume) was measured weekly. B/ The tumor volumes are indicated by black dots. The tumor growth was modelled by using a generalised logistic model. The mice were sacrificed after 40 days; at that time, the tumors were isolated and tumor weights quantified in each group (\**P* <0.05 compared with the control group). C/ Mean MIR spectra of the control group at D0 and D40 and the differential spectra of the treated groups. D/ Spectral biomarkers allowing discrimination among the groups superimposed with the mean group spectrum. E/ Tumor size prediction based on partial least square (PLS) regression showing the loadings (i.e. the correlated spectral variables and the regression model). F/ The dichotomous model to identify groups from plasma serum with, for each step, the relevant discriminant variables and the consequent confusion matrixes. G/ Metabolic pathways associated with metabolite sets that were modified between D0 and D40 for each group based on analysis of whole blood samples. H/ Heatmap analysis of metabolite levels identified at D40 showing the difference between the treated groups and the control group.

### Metabolic Mid-Infrared Signatures Predicted Tumor Size and Were Specific for Each NCLX Blocker

Because we found that curcumin and NCLX inhibitors are regulators of cancer cell metabolism, we aimed to identify some circulating biomarkers of these effects. Using MIR spectroscopy of dried plasma and serum from mice that had been xenografted with CRC cells, we looked for a specific tumor metabolic signature related to NCLX inhibitors and curcumin to gain insights into the similarities between the mechanisms of action of curcumin, CGP37157 and ITH12575 on NCLX.

The analysis was performed in control, curcumin-treated, CGP37157-treated or ITH12575-treated mice. Figure 6C presents spectral signatures between D0 and D40. The five most discriminating variables between the two groups or two subsets were identified as shown in Figure 6D: control, 3081, 2813, 1670, 984 and 902 cm^−1^; curcumin, 3048, 1738, 1730, 1520 and 1345 cm^−1^; CGP37157: 1740, 1732, 1520, 1462 and 1347 cm^−1^; and ITH12575, 3080, 1734, 1450, 1355 and 1344 cm^−1^. The signature was mainly linked to protein absorption bands at 1664, 1653, 1643, 1546 and 1534 cm^−1^. These bands are assigned to the amide vibrations of proteins, suggesting drastic changes in plasma/serum proteins. Interestingly, this signature was correlated with tumor size in the control group and was lost in the treated groups, therefore confirming a ‘treatment’ effect on tumor size (Figure 6E). To investigate whether curcumin treatment or NCLX blockers yield similar MIR signatures, we performed sequential predictive tests to determine the spectral signatures of mice at D40 according to a dichotomous decision tree (Figure 6F). Matrices easily discriminated the control group from the treated groups. Moreover, this method allowed us to discriminate the curcumin-treated group from the two other treatments (CGP37157 or ITH12575). The analysis between the CGP37157-treated and ITH12575-treated groups was trickier, which was not surprising given that ITH12575 is a derivative of CGP37157.

In summary, plasma/serum signatures were altered reproducibly through crosstalk between mice and CRC cells. The identified MIR signatures support major protein and lipid alterations in the mechanisms that underlie the anti-tumoral efficacy of curcumin and other NCLX inhibitors.

### High-Resolution Mass Spectrometry Full-Scan Analysis Revealed a Role for NCLX in the Control of Protein and Amino Acid Synthesis and Methylation

To focus on the metabolites that are modified following NCLX inhibition during tumor growth, we used LC-HRMS full-scan analysis of serum extracts from our experimental mouse model and compared metabolomic modulation of each group (between D0 and D40) (Figure 6G). The variable importance in projection (VIP) list and biochemical pathways allowed us to link or to discriminate each group with a panel of particular metabolites and pathways. For all groups, we observed a change in the metabolism of pyrimidines. Pantothenate and CoA biosynthesis pathways are affected for the control/CGP37157-treated and ITH12575-treated groups. Beta alanine metabolism is also modified under control and CGP37157 conditions. There are also specific modifications such as glycine-serine-threonine metabolism for the control group or arginine metabolism for the CGP37157-treated group. Finally, the biosynthetic pathway of unsaturated fatty acids for the ITH12575-treated group is modified. Taken together, these results suggest an important role of pyrimidine metabolism, pantothenate and CoA biosynthesis, beta alanine metabolism and glycine-serine-threonine metabolism in tumor development and that NCLX inhibitors limit the activation of these pathways. These data suggest metabolic differences over time in each group prior to implantation of tumor cells and during tumor development (Figure 6G).

To further characterise the metabolic modifications linked to NCLX through the three treatments, we compared the metabolites analysed in the treated groups to the metabolites in the control group. We found a common base of metabolite modification in our three treated group: choline, guanine, lumichrome, lyxose were up regulated (red color) and methylcytosine was down regulated (blue color) (Figure 6H). These results suggest a possible anti-tumoral effect by these pathways. We also found metabolites modification such as methionine, nepsilon, succinate or alpha glucose 1-phophate, glycerate, retinoate, deoxycytidine and trimethylamine that were only present in two groups: CGP/curcumin or CGP/ITH12575 (Figure 6H). We also found some metabolites found in only one group (Figure 6H). The combination of these results suggests an important role for NCLX in the control of protein and amino acid synthesis and methylation through the regulation of choline and possibly methionine metabolism (variation common to all 3 groups of inhibitors).

### Transcriptomic and Tissue Microarray Analysis Highlighted the Association between NCLX Expression and the MSI Status and Survival of Patients with Colorectal Cancer

Analysis of patients with CRC in the TCGA dataset showed that higher *NCLX* expression was associated with MSI status (Figure 7A). NCLX expression was next analysed on TMA-based collection of 302 patients with MSI CRC (COLOMIN cohort). We evaluated NCLX expression with regard to tumor stages. Most patients at advanced stages had lower NCLX protein expression. Of note, loss of NCLX expression was associated with tumor grade in poorly differentiated cancers (Figure 7B, a–d; *P* < 0.0001). Subsequent analysis revealed that the NCLX expression level was inversely associated with perineural invasion (*P* < 0.0001) and vascular emboli (*P* < 0.02), suggesting a protective role of this exchanger in CRC. We found that NCLX was expressed in 237 of 261 normal tissue samples (91%), with granular cytoplasmic staining (Figure 7C). In the cancerous tissues analysed with TMA, NCLX remained expressed in 183 of 321 samples (57%), with either moderate-intensity (Figure 7C, panel NCLX+) or high-intensity (Figure 7C, panel NCLX++) staining; in 138 cancer tissue samples, NCLX expression was lost (Figure 7C, panel NCLX−).

**Figure 7:**
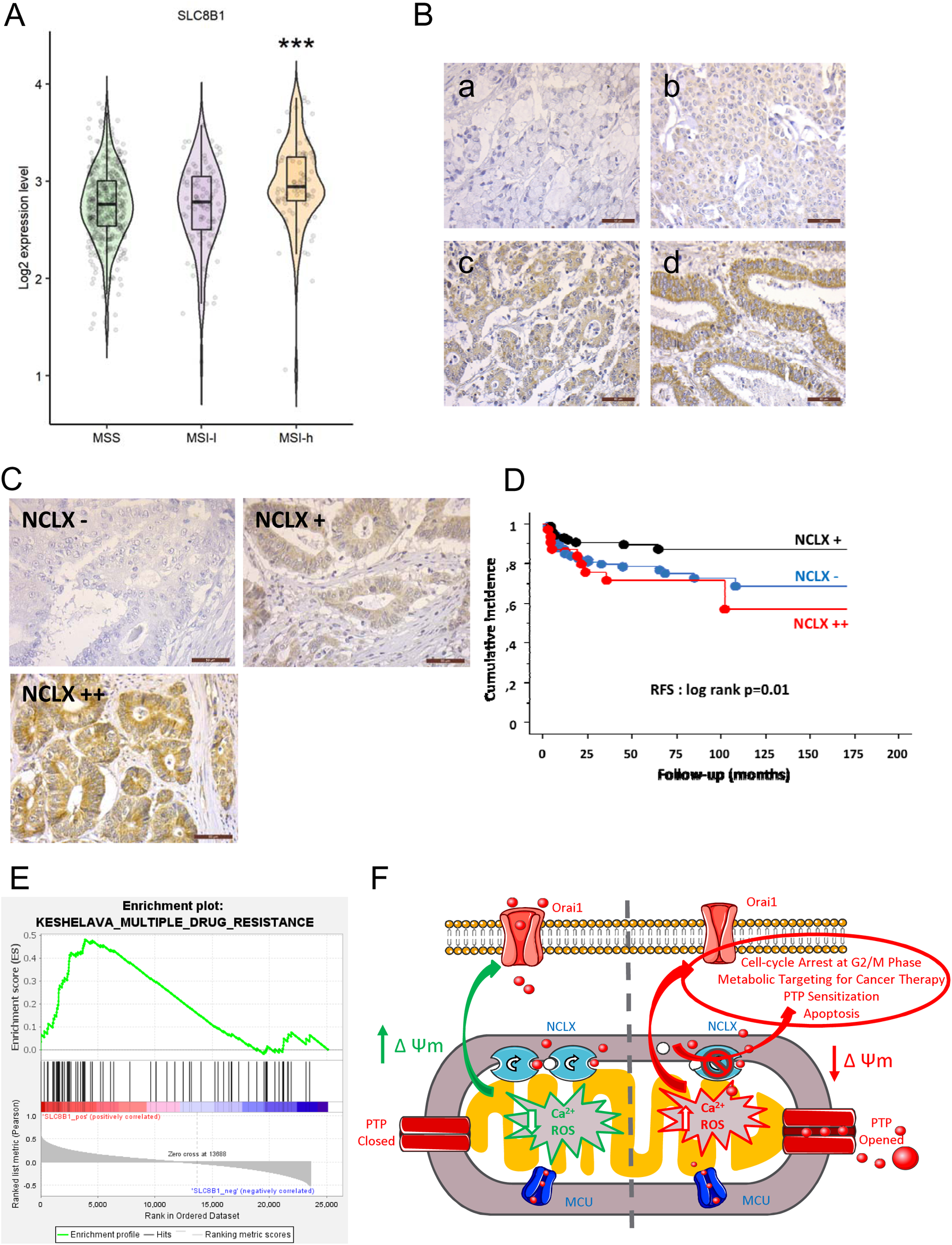
NCLX expression and clinical parameters of a cohort of patients with MSI CRC. A/ Boxplots showing the mRNA expression profile of NCLX in primary CRC tissues from the TCGA dataset according to their MSI/MSS status (MSS, n = 423; MSI-Low [MSI-L], n = 100; MSI-High [MSI-H], n = 81). B/ Representative NCLX expression during the differentiation of the colonic epithelium (COLOMIN cohort). C/ Representative NCLX immunostainings of three groups of patients with CRC classified according to their NCLX expression level (COLOMIN cohort). D/ Kaplan–Meier plots (cumulative incidence in function of follow-up of patients with CRC) showing RFS of the COLOMIN cohort (*P* = 0.01 by the log-rank test). E/ GSEA of the KESHELAVA multiple drug resistance signature in primary MSI CRC tissues from the TCGA dataset.

Based on Cox univariate analysis, NCLX expression was associated with RFS (Figure 7D). Tumors with loss of NCLX expression (Figure 7C, panel NCLX-) recurred more than those that retained NCLX expression (Figure 7C, panel NCLX+). Surprisingly, tumors with increased NCLX expression (Figure 7C, panel NCLX++) had an even worse prognosis than the patients with retained or lost NCLX expression. This finding suggests that NCLX is normally expressed normally in non-cancerous epithelial cells and its loss of expression is associated with an aggressive progression of epithelial tumors. At the same time, however, overexpression of this exchanger can lead to a worsened evolution of the disease. In line with these data, GSEA of the transcriptomic CRC dataset from TCGA revealed a positive correlation between *NCLX* expression and the multiple drug resistance signature in patients with MSI (Figure 7E). Our main results are summarised in a schematic diagram (Figure 7F).

## Discussion

Understanding the mechanisms of curcumin-mediated apoptosis and subsequent cell cycle arrest has important implications for anti-cancer therapy (27). Over the last few years, the mechanisms of Ca^2+^ signaling has been greatly improved, especially how its alterations in cancer participate in modifying key processes such as proliferation, metabolism and sensitivity to cell death (18). Recent evidences highlighted that mtCa^2+^ is closely associated with the hallmarks of cancer and MCU and NCLX play key roles in CRC carcinogenesis (19)(28). However, little is known about the role of curcumin on mtCa^2+^ fluxes. The main results of the present study are the demonstration of the mtCa^2+^-dependent effects of curcumin on NCLX and the potential anti-tumoral effect of NCLX blockers in CRC. Ca^2+^ directly regulates the expression and activation of cell cycle protein complexes (29). NCLX silencing or curcumin treatment led to an accumulation of cells at the G2/M checkpoint.

Curcumin induced a high mtCa^2+^ level by blocking mtCa^2+^ efflux through NCLX. Our data showed that pharmacological inhibition of NCLX in CRC cells caused mtCa^2+^ overload and an increase in mitochondrial and cytosolic ROS production associated with a mitochondrial membrane depolarization. Kostic et al., demonstrated that mild fluctuations in ΔΨm, which do not affect Ca2+ influx, are sufficient to strongly regulate NCLX (30). Furthermore complete mitochondrial depolarization and deenergization leads to reversal of NCLX activity preceding toxic mtCa^2+^ accumulation (31). Our results do not allow us to show whether curcumin directly blocks NCLX or whether the action is governed by an effect on the mitochondria membrane potential. However, the study from Mustapha *et al*. showed that the increase of mitochondrial calcium is nearly occurring in parallel with curcumin accumulation, and precedes ROS production and the ΔΨm decreasing, suggesting a potential effect of curcumin on NCLX in the first instance (32)(33).

Intrinsic apoptosis involves mtCa^2+^ and mtROS overload and mitochondria membrane depolarization leading to mPTP opening and release of cytochrome *c* (15). Curcumin treatment induced apoptosis in a dose-dependent manner. We found that curcumin treatment can lead to a sustain increase of the rate of mPTP activation in CRC cells. Consistently, NCLX KO cells and cells treated with CGP37157 had greater mPTP activation. These effects were reversed by pre-treatment with CsA. These effects of curcumin on mtCa^2+^-triggered apoptosis have already been described. Indeed, the researchers showed that ruthenium red (RR), an inhibitor of MCU, specifically inhibits curcumin-induced apoptosis in U937 cells (34). Similarly, curcumin-induced increases in mtCa^2+^ were markedly inhibited by pretreatment with RR or Ru360 in MDA-MB 435S cancer cells (33). In this study, authors investigated the role of Ca^2+^ in curcumin-induced <u>paraptosis</u>. Curcumin induced mitochondrial Ca^2+^ overload selectively in the malignant breast cancer cells (33).

The authors found that CGP-37157 and bortezomib co-treatment activates paraptotic signals in cancer cells, similar to curcumin. The cell death induced by CGP-37157 plus bortezomib or curcumin were dose-dependently inhibited by cycloheximide (CHX, a protein synthesis blocker), N-Acetyl-Cysteine (NAC) (an antioxidant), U0126 (a MEK inhibitor), or SP600125 (a JNK inhibitor) suggesting that the underlying mechanism of the cell death by CGP-37157 plus bortezomib may be very similar to that of curcumin-induced paraptosis (33).

SOCE activates a mitochondrial redox transient which is dependent on NCLX and is required for preventing Orai1 inactivation through oxidation of a critical cysteine (Cys195) in the third transmembrane helix of Orai1 (22). In agreement with this the replacement of the reactive cysteine residue with serine at a specific Orai1 (C195S) site significantly altered the sensitivity of I_CRAC_ to curcumin, suggesting that the Cys195 is the target residue for the inhibitory actions of this compounds (35). As we previously showed in CRC cell lines with NCLX KO (19) the pharmacological blockade of NCLX by curcumin or CGP37157 decrease SOCE in CRC cells. We also confirmed the effect of curcumin treatment on Orai1-dependent CRAC current.

The fine regulation of Ca^2+^ in MAM is paramount for the management of metabolism and cell death (36). Cancer cells need a constitutive transfer of Ca^2+^ from the ER to mitochondria for their survival (37)(38). Our data showed that NCLX targeting disrupted MAM contacts, a phenomenon that may be involved in the activation of cell death and metabolic reprogramming. Remodeling of cancer cell metabolism according to nutrient availabilities is a hallmark of cancer (36). The inhibition of constitutive mtCa^2+^ uptake induced a bio-energetic crisis that resulted in metabolic reprogramming. However, the effect of mtCa^2+^ efflux has been poorly described. Recently, Assali *et al*. highlighted a key pathway through which mtCa^2+^ efflux permits a robust activation of respiration in response to profound mitochondrial respiratory uncoupling, while preventing the activation of cell death pathways (39).

In our previous study, we reported that NCLX loss in CRC cells leads to reduced oxygen consumption, increased glycolysis and increased transcription of major glycolytic enzymes and transporters (19). This increased transcription of glycolytic enzymes is under the control of HIF1α. This transcriptional factor is upregulated in response to increased mtROS resulting from loss of NCLX and subsequent increase in mtCa^2+^. In the present study, we demonstrated that curcumin and NCLX inhibition suppressed mitochondrial respiration, amino acid metabolism and HIF1α-dependent reduced consumption of glycolysis substrates. The latter finding is in contrast to the effect observed by Pathak et al., suggesting either an adaptation (metabolic reprogramming) of siNCLX-treated cells and NCLX KO cells, or an off-target effect of curcumin or CGP37157. Additional experiments should be carried out to refine the understanding of the mechanism of action of NCLX blockers.

*In vivo*, we demonstrated that NCLX inhibitors reduced the tumor size of NMRI nude mice grafted with HCT116 cells. The effect of curcumin on tumor growth may be explained by its effect on cell proliferation (see Figure 1) and by its anti-angiogenic potency as observed in Supplementary Figure 5 (showing a completely inhibited endothelial tube formation). We thus reasoned that cancer cells with or without NCLX blockers could affect the global metabolism of mice. Interestingly, using MIR spectroscopy on whole blood from xenografted mice, we found a five-element signature in the control group that correlated with tumor size. This correlation was lost in the treated groups, suggesting the possibility to follow and to model tumor growth with minimally invasive peripheral biomarkers. Moreover, spectral analysis at D40 allowed us to isolate a five-element signature specific for each group, thereby allowing discrimination of the experimental groups. Intriguingly, CGP37157 and curcumin caused identical spectral modifications at 1730–1740, 1520 and 1345–1347 cm^−1^, suggesting a similar global effect in these particular subgroups. To further support our findings, we used metabolomics to identify shared pathways and metabolites among treatments compared with the control group; this analysis produced a similar pattern as seen with MIR spectroscopy. Metabolites of methylcytosine, choline and guanine appeared to be particularly affected by the blockade of NCLX. These results showed that NCLX is a key ‘crossroad’ factor in the control of protein and amino acid synthesis and methylation. Choline modulates epigenetic marking of genes, and choline deficiency is correlated with the silencing of several tumor suppressor genes responsible for DNA repair (hMLH1) and cell cycle regulation (p15 and p16)(40). In the future, we aim to distinguish unique metabolic profiles or easy-to-implement signatures that would be useful as minimally invasive biomarkers of NCLX inhibition in a clinical context.

In the first randomised controlled trial using curcumin in combination with FOLFOX chemotherapy for patients with metastatic CRC, researchers showed that this combination is a safe and tolerable treatment that could benefit patients with longer overall survival, but the number of patients was too low (n = 27) to make definitive conclusion (41). MSI-H CRC tumors have distinct clinical and pathological features such as frequent early stage, mucinous histology, poor differentiation and proximal location. In early-stage CRC, MSI can select a group of tumors with a better prognosis due to high immune infiltration; on the other hand, in metastatic disease it seems to confer a negative prognosis due to immune escape. Although there are numerous conflicting results, a large body of preclinical and clinical data suggests these tumors might be resistant to 5-FU, especially at stage II. In our cohort of patients with MSI CRC, NCLX expression decreased in advanced CRC stages. NCLX expression was associated with poor tumor differentiation but lower perineural sheathing and/or vascular emboli. Loss of NCLX expression was associated with worse RFS in Cox univariate analysis. Surprisingly, we found a subgroup of patients with MSI CRC (13%–15%) with increased NCLX expression; they had an even worse prognosis than patients who had maintained or lost NCLX expression. Additional investigations will be necessary to understand the origin of this subgroup with increased NCLX expression. Moreover, CGP37157 has been reported to prevent neurotoxicity in response to the chemotherapeutic agent salinomycin. This protective effect was not observed when head and neck squamous cell carcinoma cells were co-treated with salinomycin and CGP37157 (42). Prospective trials need to evaluate the combination of chemotherapy plus curcumin in patients with NCLX-expressing MSI CRC to improve the efficacy and decrease the adverse events of standard treatment.

In summary, our findings, obtained from CRC cell lines, tumors and circulating factors from a model of xenografted NMRI nude mice, dataset analysis and TMA from human MSI CRC tumors provide very strong evidence that curcumin and NCLX blockers inhibit CRC cell growth by enhancing cellular oxidative stress and mtCa^2+^ overload associated to a mitochondria membrane depolarization, thereby leading to mPTP-dependent cell apoptosis. Our works also show that reducing mitochondrial respiration, HIF1α and MAM perturbations contribute to these mechanisms. These findings support a novel anti-tumoral mechanism for NCLX, particularly in patients with MSI CRC, a clinical finding that has never been described before to our knowledge.

## Materials and Methods

### Study Design, Patient Characteristics and Survival Analysis

From 2003 to 2015, all patients with MSI CRC classified by molecular testing from the cancer biology departments of Poitiers and Tours University Hospitals were included in the multi-centre prospective COLOMIN cohort (*Cohorte nationale des cancers colorectaux avec instabilité microsatellitaire*). This study was designed in accordance with legal requirements and the Declaration of Helsinki and was approved by the ethics committee of CHU Poitiers and CHRU Tours, France (Comité de protection des personnes Ouest III, n°DC-2008-565 and n°2018-039).

The main patient characteristics (gender and age) and tumor characteristics (tumor site, Tumor Node Metastasis [TNM] stage, VELIPI criteria [vascular emboli, lymphatic invasion, or perineural invasion], tumor grade, tumor perforation and initial bowel obstruction) were collected. Whether a case was germline (Lynch syndrome) or sporadic dMMR/MSI was determined as described previously (4). Information regarding disease recurrences and deaths were collected.

Tumor DNA was extracted from all tumors by using the KAPA Express Extract Kit (Roche, Basel, Switzerland). The MSI phenotype was assessed by analysing microsatellite loci comprising five mononucleotide markers: BAT-25, BAT-26, NR21, NR24 and NR27 (Pentaplex panel) (4). MSI was defined by the presence of instability affecting at least three of the five markers. In the case of one or two markers with instability, a comparative analysis of normal colon and tumor DNA was performed.

RFS was defined as the time from the date of primary tumor curative surgery until the date of recurrence. Patients without recurrence were censored at the last follow-up. Kaplan–Meier survival curves were generated and groups were compared using the log-rank test. *P* < 0.05 was considered statistically significant. Survival analyses were performed in the Statview environment.

### Tissue Microarray and Immunohistochemistry

Tissue microarray (TMA) construction: TMAs were constructed using formalin-fixed paraffin-embedded tissue samples. For each case, nine cores of 0.6-mm diameter (three in the tumor centre, three in the invasive front and three in non-tumor tissue) were transferred from the selected areas to the recipient block, using a TMA workstation (Manual Tissue Arrayer MTA Booster, Alphelys, France). Serial 3-µm sections of the TMA blocks were used for immunohistochemistry. Every tenth section was stained with haematoxylin and eosin to check that the cores adequately represented diagnostic areas.

Immunohistochemistry: Slides were deparaffinised, rehydrated and heated in citrate buffer (pH 6) for antigen retrieval. After blocking endogenous peroxidase with 3% hydrogen peroxide, the primary antibodies were incubated. After incubation with the primary antibody NCLX (HPA040668, Sigma-Aldrich, Saint-Quentin-Fallavier, France) diluted 1:50 overnight at 4°C, immunohistochemistry was performed by using the streptavidin-biotin-peroxidase method with diaminobenzidine as the chromogen (Kit LSAB, Dakocytomation, Glostrup, Denmark). Negative controls were obtained after omission of the primary antibody or incubation with an irrelevant antibody.

Scoring of antibody staining: A semi quantitative score was assigned based on the intensity of staining: 0 (no staining), + (moderate staining) and ++ (strong staining).

### Cell Culture, Transfection and Treatments

HCT116 (RRID:CVCL_0291), LoVo (RRID:CVCL_0399) and HT-29 (RRID:CVCL_0320) SW48 (RRID:CVCL_1724) cell lines were purchased from ATCC (Manassas, VA, USA) and cultured in McCoy’s medium (Gibco, Thermo Fisher, Illkirch, France) supplemented with 10% foetal bovine serum (FBS, Eurobio, Les Ulis, France), without antibiotics. NCM356 cells (Incell Corporation, LLC, San Antonio, TX, USA) were cultured in high-glucose Dulbecco’s Modified Eagle Medium (DMEM) (Sigma-Aldrich) supplemented with 10% FBS and antibiotics. The cell lines were cultivated at 37°C in a humidified incubator with 5% CO_2_. The cell lines were maintained and *Mycoplasma* absence was tested regularly using the PlasmoTest kit (HEK blue, Invivogen, Toulouse, France). HCT116 NCLX KO cells are the same cells described in Pathak et al., 2020 (19). NCLX KO cells were generated using a guide RNA (g1) which resulted in a single cut at nucleotide 150 in exon one causing a frameshift mutation and introduction of a stop codon at position 180 in the NCLX open reading frame.

### Transcriptomic Analysis

The Cancer Genome Atlas (TCGA) dataset of patients with CRC was downloaded from cbioportal^47^ and analysed as described previously (43) (44). Briefly, clinical and transcriptomic data were acquired using the *gdsr* package in the R environment. Microsatellite instability (MSI) and microsatellite stable (MSS) annotations were downloaded from the Broad Institute’s firehose repository (https://gdac.broadinstitute.org/).

Gene Set Enrichment Analysis (GSEA) was performed on patients with MSI-h CRC from the TCGA transcriptomic dataset using the GSEA JAVA application from the Broad Institute with default parameters (http://software.broadinstitute.org/gsea).

### Compounds

Curcumin (10 µM) was purchased from TOCRIS (Bristol, UK); CGP37157 (10 µM) was purchased from Sigma-Aldrich (Merck, Darmstadt, Germany); ITH12575 (10 µM) was purchased from Sigma-Aldrich.

### Transfection

Briefly, 2.5 × 10^5^ cells/well were plated in 6-well plates in McCoy’s medium supplemented with 10% FBS without antibiotics. The cells were incubated with a mix of small interfering RNA (siRNA) and lipofectamine in medium without serum for 6 h. After incubation, an equal volume of medium with serum was added to each well. The siRNA sequences directed against NCLX were purchased from Dharmacon (L-007332-00-0005) (La Fayette, CO, USA).Measurement of NCLX WT and NCLX S468T on cell viability in HCT116 cells was achieved by transient overexpression of NCLX WT and NCLX S468T ((45)) as previously described (45). Briefly, 1 × 106 cells were incubated with a solution containing 1.5 μ of NCLX WT or NCLX S468Tand 3 μl Lipofectamine 2000 for 3 h. One day later, transfected cells were transferred and treated with 10 µM curcumin for 24h in 24-well plates.

### Reverse Transcription and Real-Time Polymerase Chain Reaction

Total RNA was collected using the Nucleospin RNA Kit (Macherey-Nagel, Hoerdt,France) and transcribed into complementary DNA (cDNA) with the GoScript Reverse Transcription System (Promega, Madison, WI, USA). cDNA was then amplified with the SYBR Green Master kit (Roche, Mannheim, Germany) using a Light Cycler 480 apparatus. PCR was performed in 40 cycles of 15 s at 95°C and 45 s at 60°C. The average Δ t value was calculated for each cell line with respect to the housekeeping gene *HPRT1*. The primers sequences used are:

- HPRT1: forward 5′ CAT-TAT-GCT-GAG-GAT-TTG-GAA-AGG 3′ reverse 5′ CTT-GAG-CAC-ACA-GAG-GGC-TAC-A 3′;
- NCLX: forward 5′ GCC-AGC-ATT-TGT-GTC-CAT-TT 3′ reverse 5′ T-TCG-TCT-CGG-CCA-CTT-AC 3′;
- MCU: forward 5′ CGC-CAG-GAA-TAT-GTT-TAT-CCA 3′ reverse 5′ CTT-GTA-ATG-GGT-CTC-TCA-GTC-TGT-T 3′
- MICU1: forward 5′GAG-GCA-GCT-CAA-GAA-GCA-CT 3′ reverse 5′ CAA-ACA-CCA-CAT-CAC-ACA-CG 3′
- MICU2: forward 5 GGC-AGT-TTT-ACA-GTC-TCC-GC 3′, reverse 5′ AAG-AGG-AAG-TCT-CGT-GGT-GTC 3′
- STIM1: forward 5′ GCC-CTC-AAC-ATA-GAC-CCC-AG 3, reverse 5′ TCC-ATG-TCA-TCC-ACG-TCG-TCA 3′
- STIM2 forward 5′ TTG-GAC-CCT-TGA-AGA-CAC-TCT 3′ reverse 5′ CCA-GTT-ATG-AGG-TGG-GCG-TG 3′
- Orai1 5′ AGG-TGA-TGA-GCC-TCA-ACG-AG 3′, reverse 5′ CTG-ATC-ATG-AGC-GCA-AAC-AG 3′

TRPC1 5′ TTA-CTT-GCA-CAA-GCC-CGG-AA 3′, reverse 5′ CTG-CTG-GCA-GTT-AGA-CTG-GG 3′

### Semi-Quantitative Real-Time Polymerase Chain Reaction Analyses (Redox Profiling)

Semi-quantitative real-time polymerase chain reaction (RT-qPCR) analyses were performed as described previously by Kouzi et al (46). Briefly, HCT116 cell line was treated for 24 h with 10 µM curcumin and RNA was isolated using the Maxwell® 16 IVD system and the Maxwell Simply RNA Tissue Kit according to the manufacturer’s instructions (Promega). RNA was reverse transcribed using the SuperScript® VILO cDNA Synthesis Kit (Invitrogen, Waltham, MA, USA) and RT-qPCR was performed using the Universal Probe Library System (Roche, Penzberg, Germany). cDNA, primers, probes and TaqMan master mix were mixed and PCR was run in a LightCycler® 480 (Roche). The mean cycle threshold (Ct) of the human *ACTB* and *RPL13A* genes were used as endogenous controls to normalise the expression of target genes. Each reaction was performed in triplicate. Relative quantification (RQ) gene expression was evaluated by the 2^−Ct^ method (47). The list of primers is provided in Supplementary Table S1.

### Flow Cytometry Analysis

#### Cell Cycle

Cells were harvested by using trypsin-ethylenediaminetetraacetic acid (EDTA, Thermo Fisher Scientific, San Jose, CA, USA) 24 h after treatments, washed in 1X phosphate-buffered saline (PBS), fixed in cold 70% ethanol and incubated at −20°C for at least 2 h. Subsequently, the samples were resuspended in PBS with RNase and stained with 0.25 mg/mL propidium iodide (PI) for 30 min in the dark at room temperature. The DNA content of stained cells was analysed using a Gallios flow cytometer (Beckman Coulter, Villepinte, France). For each sample, a minimum of 5 × 10^4^ cells were evaluated. Analyses were done using the Kaluza 1.3 software (Beckman Coulter).

#### Apoptosis

Cells were harvested by Accutase (Thermo Fisher Scientific, San Jose, CA, USA), washed with 1X PBS and then resuspended in annexin binding buffer following the manufacturer’s instructions (Annexin-PI kit, Thermo Fisher Scientific, San Jose, CA, USA) and analysed by flow cytometry as described above.

### Spheroid Cell Viability Test

The spheroid cell viability test was performed according to the manufacturer’s conditions (Cultrex proliferation – cell viability 3510096K, Bio-Techne, Abingdon, UK). The cells were cultured in extracellular matrix supplemented with Epidermal Growth Factor (EGF) in 96-well plates with round bottoms for 48 h and then treated with 10 µM curcumin for 72 h. To capture the entire spheroid, image fields starting at the centre of the well were collected using a 20× objective lens (Nikon, Champigny-sur-Marne, France).

### 3-(4,5-Dimethylthiazol-2-yl)-2,5-Diphenyltetrazolium Bromide (MTT) Assay

Cells were cultured and treated with 1–10 µM curcumin for 24, 48 or 72 h. 3-(4,5-Dimethylthiazol-2-yl)-2,5-diphenyltetrazolium bromide (MTT) solution (0.5 mg/ml) was added to each well prior to incubation. The cells were incubated at 37°C for 45 min. Afterwards, the supernatant was removed and dimethyl sulphoxide (DMSO) was added to dissolve formazan crystals and absorbance was read at 570 nm using a Mithras LB 940 Multimode Microplate Reader (Berthold, Bad Wildbad, Germany).

### Sulforhodamine B (SRB) Assay

Cells were cultured and treated with 1–10 µM curcumin for 24, 48 or 72 h Cell viability, survival in HBSS and short-term toxicity were evaluated using standard sulforhodamine B (SRB) method after 72h, 48 h and 24 h treatments, respectively. Briefly, cells were fixed with 50% trichloroacetic acid for 1 h at 4 °C and stained for 15 min with 0.4% SRB solution. Cells were washed 3 times with 1% acetic acid and dye was dissolved with 10 mM trisbase solution over 10 min. Absorbance at 540 nm was read using BioTek Spectrophotometer.

### NanoLuc activity assay

The HIF1α NanoLuc protein-reporter HCT116 and HCT116 NFE2L2-C-terminal Luciferase cell lines were purchased from Horizon Discovery (Cambridge, UK). Cells were seeded in 96-well plates at 3 × 10^4^ cells/well and incubated for 24 h in normoxic or hypoxic conditions (1% O_2_) with or without 10 µM curcumin or 10 µM CGP37157. Luciferase activity was measured using the Nano-Glo® Reagent (Nano-Glo® Luciferase Assay, Promega) following the supplier’s instruction. Luminescence intensity values were measured by FlexStation-3 (Molecular Devices, San José, CA, USA).

### Store-Operated Calcium Entry Measurement by Fura-2 AM

Cells were plated in 96-well plates at 2 × 10^4^ cells/well 24 h before the experiment. Adherent cells were for loaded with the ratiometric dye Fura2-acetoxymethyl ester (AM; 5 μM) for 45 min at 37°C then washed with PBS supplemented with Ca^2+^. During the experiment, the cells were incubated with Ca^2+^-free physiologic saline solution (PSS) solution and treated with 2 µM thapsigargin (TG, T7458, Life Technologies, Thermo Fisher) to deplete intracellular store of Ca^2+^. Ca^2+^ entry was stimulated by injection of 2 mM of CaCl_2_. Fluorescence emission was measured at 510 nm using the FlexStation-3 (Molecular Devices) with excitation at 340 and 380 nm. The maximum fluorescence (peak of Ca^2+^ influx [F340/F380]) was measured and compared with the normal condition. NCLX was inhibited by 10 µM CGP37157 or 10 µM curcumin.

### Mitochondrial Ca^2+^ Measurements

To measure mtCa^2+^ using the Rhod-2 AM dye (543 nm/580–650 nm), the cells were cultured at 60%–80% confluency in 96-well plates. The cells were washed with medium without FBS and antibiotic-antimycotic agents. Then, the cells were incubated in medium containing 3 µM Rhod-2 AM (without FBS and antibiotic-antimycotic agents) at 37°C for 45 min. The cells were washed and kept in PBS (HEPES-buffered saline solution, 140 mM NaCl, 1.13 mM MgCl_2_, 4.7 mM KCl, 2 mM CaCl_2_, 10 mM D-glucose, and 10 mM HEPES, adjusted to pH 7.4 with NaOH) containing 2 mM CaCl_2_ for imaging. The cells were stimulated with 100 µM adenosine triphosphate (ATP) in PBS containing 2 mM CaCl_2_. The timelapse images were acquired using the FlexStation-3 (Molecular Devices). To measure mtCa^2+^ using the mt-Cepia, Cultured cells were transfected using Lipofectamine 2000 (Invitrogen) with 1,5µg pCMV CEPIA3mt 2 days before imaging. 24hours before the experiments transfected cells were plated in 96-well plates. The cells were stimulated with 100 µM adenosine triphosphate (ATP) in PBS containing 2 mM CaCl_2_. The timelapse images were acquired using the FlexStation-3 (Molecular Devices). *N*-methyl-d-glucamine (NMDG) was purchased from Sigma-Aldrich. PCMV CEPIA3mt was a gift from Masamitsu Lino (Addgene plasmid # 58219; http://n2t.net/addgene:58219; RRID:Addgene_58219).

Calcium Green-5N (Invitrogen) on permeabilized cells beforehand with digitonin was used to measure mitochondrial Ca^2+^ uptake and release experiments.

HCT116 permeabilized cells suspended in intracellular Na^+^-free buffer (130 mM KCl, 10 mM Tris-MOPS (pH 7.4), 10 μM EGTA-Tris, 1 mM KPi, 5 mM malate, 5 mM glutamic acid, 1 μM Ca^2+^ green-5N)..At the starting point Curcumin or CGP37157 were added following by one pulse of 10µM Ca^2+^ at 2min. Next, 0.5 μM Rutenium Red (RR), an inhibitor of MCU, was added at 6min to block Ca^2+^ uptake. Next, the Na^+^-induced Ca^2+^ release was initiated by the addition of a Na^+^ pulse (10mM at 7min). Ca^2+^ uptake/release measurements were performed at 25°C on a F2710 spectrofluorometer (Hitachi) with the following parameters, λ_ex_ = 505 nm, λ_em_ = 535 nm, slit width: 2.5 nm.

### Mitochondrial and Cytoplasmic Reactive Oxygen Species Measurements

Cells were cultured in 6-well plates at 70%–90% confluency. To measure mitochondrial reactive oxygen species (mtROS), 1 × 10^6^ cells were stained with 5 µM MitoSOX (Molecular Probes, Thermo Fisher) in PBS at 37°C for 40 min. The intensity of staining was measured using a Gallios flow cytometer (Beckman Coulter) and analysed with the FlowJo software (Tree Star, Ashland, OR, USA).

Cytoplasmic ROS production was measured by using the 2,7′ lorofluorescein diacetate (DCFDA) assay according to manufacturer’s protocol (Molecular Probes, Thermo Fisher). Cells were seeded at 2 × 10^3^ cells/well of 96-well plates 24 h before the experiment. The cells were washed with PSS containing 2 mM Ca^2+^ and then treated with PSS containing 2 mM Ca^2+^ and 10 µM DCFDA with or without curcumin or CGP37157. The fluorescence emission was measured at 520 nm using the FlexStation-3 (excitation: 500 nm). The fluorescence rate is reported as relative fluorescence units (RFU) normalised to time 0.

### Measurement of Mitochondrial Permeability Transition Pore Opening

Direct assessment of mitochondrial permeability transition pore (mPTP) opening in HCT116 cells was performed by loading cells with calcein-AM (Sigma-Aldrich) and CoCl_2_, resulting in mitochondrial localisation of calcein fluorescence. Specifically, the cells were loaded with 1 μM calcein-AM for 30 min at 37°C in 1 mL (PSS 2 mM Ca^2+^, pH 7.4) with 1 mM CoCl_2_. Subsequently, they were incubated with medium free of calcein-AM and CoCl_2_ and then incubated in reoxygenation medium. mPTP opening was determined from the reduction in mitochondrial calcein signal (slope value) after injection of ionomycin (5 μ). Fluorescence emission was measured at 515 nm using the FlexStation-3. Calcium Green-5N (Invitrogen) on permeabilized cells beforehand with digitonin was used to measure the Ca^2+^ retention capacity that reflects mitochondrial function. Ca^2+^ uptake was followed by measuring extra-mitochondrial Calcium Green-5N until mPTP opening was achieved. Fluorescence emission was measured at 530 nm.

### Measurement of Mitochondrial Potential

Cells were cultured in 96-well plates at 50–60% confluency. The next day, cells were loaded with Tetramethyl rhodamine (TMRE) dye. The cells were stained with 100 nM TMRE dye in complete growth media and kept at 37°C in 5% CO_2_ for 20–30 min. CCCP (50 µM) was used as a positive control.. The intensity of staining was measured and analyzed on FlexStation-3 (Molecular Devices).

### Patch Clamp Recording

Traditional patch clamp electrophysiological recordings were carried out using an Axopatch 200B and Digidata 1440A (Molecular Devices, San José, CA, USA). Pipettes were pulled from borosilicate glass capillaries (World Precision Instruments, Sarasota, FL,USA) with a P-1000 Flaming/Brown micropipette puller (Sutter Instrument Company, Novato, CA, USA) and polished using DMF1000 (World Precision Instruments). The resistance of filled pipettes was 2–4 MΩ. Under the whole-cell configuration, only cells with < 8 MΩ series resistance and tight seals (> 16 GΩ) were chosen for recordings. The cells were maintained at a +30 mV holding potential during experiments. The Clampfit 10.1 software was used for data analysis. The solutions used for this procedure were:

- Bath solution: 115 mM sodium methanesulfonate, 10 mM CsCl, 1.2 mM MgSO_4_, 10 mM HEPES (4-(2-hydroxyethyl)-1-piperazineethanesulfonic acid), 20 mM CaCl_2_ and 10 mM glucose (pH adjusted to 7.4 with NaOH);
- Pipette solution: 115 mM caesium methanesulfonate, 10 mM caesium BAPTA (Tetra-sodium 1,2-bis(2-aminophenoxy)ethane-N,N,N′,N′-tetraacetate), 5 mM CaCl_2_, 8 mM MgCl_2_ and 10 mM HEPES (pH adjusted to 7.2 with CsOH). Based on the Maxchelator software (http://maxchelator.stanford.edu/), the calculated free Ca^2+^ was 150 nM.

Calcium release-activated channel (CRAC) currents were recorded in the ORAI1 knockout (KO) HEK293 cells expressing 4 µg EYFP-STIM1 along with 1 µg Orai1-CFP. The transfected cells were incubated with or without 10 µM curcumin for 20 min before patch clamp recording.

### Identifying Mitochondria–Endoplasmic Reticulum Contact Sites by using Bioluminescence Resonance Energy Transfer

Bioluminescence resonance energy transfer (BRET) assays were carried out as described previously (25). Briefly, 5–10 × 10^3^ HCT116 cells containing the BRET-based biosensor named MERLIN (Mitochondria-ER Length Indicative Nanosensor) were seeded into a white 96-well plate (Greiner Bio-One, Kremsmünster, Austria) for 48 h. Next, cells were incubated with CGDP for 3 h or transfected with control/NCLX small interfering RNA (siRNA) (30–50 nM) for 48 h. Then, cells were incubated with 5 μM coelenterazine h (Promega, Madison, WI, USA) in phosphate-buffered saline (PBS) for 5 min in the dark. Subsequently, BRET1 measurements were carried out in an INFINITE M PLEX plate reader (Tecan, Switzerland) at room temperature, measuring *Renilla* luciferase 8 (Rluc8) and mVenus emissions. In every experiment, Rluc8-L1-B33C (donor of the BRET pair) alone was transfected in HCT 116 cells 16 h prior to BRET measurement. The BRET signal was calculated as the acceptor emission relative to the donor emission and corrected by subtracting the background acceptor/donor ratio value detected when RLuc8-L1-B33C is expressed alone.

### SeaHorse Analysis

The cellular oxygen consumption rate (OCR) and extracellular acidification rate (ECAR) data were obtained using a Seahorse™ XF96 Flux Analyzer from Seahorse Bioscience (Agilent Technologies, Santa Clara, CA, USA) as we described previously (48). The experiments were performed according to the manufacturer’s instructions. Briefly, HCT116 cells were seeded in XF96 cell culture plates at 2 × 10^4^ cells/well, then cells were treated with 10 µM curcumin or CPG37157 for 12 h. On the day of analysis, the culture medium was replaced with XF Dulbecco’s Modified Eagle Medium (DMEM, Thermo Fisher Scientific, San Jose, CA, USA) supplemented with 2 mM glutamine and lacking bicarbonate (pH 7.4). The cells were then incubated at 37°C in a non-CO_2_ incubator for 1 h and measurements were performed as described in the relevant figure legends. Sequential injection of 10 mM glucose, 1 µM oligomycin, 100 µM dinitrophenol (DNP) and 0.5 µM rotenone/antimycin A permitted the determination of the main respiratory and glycolytic parameters. Finally, the data were normalised to the amount of DNA present in the cells and assayed using Cyquant® Cell Proliferation Assay kit (Thermo Fisher Scientific, San Jose, CA, USA). The data were acquired with the Seahorse Wave Controller and analysed with the Seahorse Wave Desktop Software.

### OmniLog Analysis

Metabolic profiling was studied by using the OmniLog® Phenotype Microarray™ system (Biolog, Hayward, CA, USA) to evaluate the cell’s ability to metabolise 367 different carbon and energy substrates (46). According to the manufacturer’s instructions, HCT116 cells were cultured at 2 × 10^4^ cells/well for 24 h in the presence of 10 µM curcumin in PM-M1, PM-M2, PM-M3 and PM-M4 plates (Biolog) in a phenol red-free RPMI-1640-based medium depleted of carbon energy sources (IFM1 medium, Biolog) supplemented with 0.3 mM glutamine, 5% fetal calf serum, 100 U/mL penicillin and 100 μL streptomycin.

Following the incubation, 10 µL Redox Dye Mix MB (Biolog, Hayward, CA, USA) was added. Kinetically, tetrazolium reduction was measured at 37°C over 6 h with the OmniLog® automated incubator-reader. Data are acquired with the Data Collection 2.4 software and analysed with the PM-M Kinetic and PM-M Parametric software programs. The area under the curve (AUC) of each metabolite consumption was determined using the *opm* package in the R environment after correction of the background signal with the negative control wells.

### Subcutaneous Colorectal Cancer Xenograft

CRC xenografts were established by injecting 5 × 10^6^ tumor cells per mouse. Swiss nude mice were randomised regarding their tumoral volume into treatment groups (n = 8–9/group) after the mean tumor volume reached ~200 mm. Mice were given USP saline with vehicle(DMSO/EtOH) MWF, curcumin at 150 mg/kg per dose (MWF) or CGP37157 at 150 mg/kg (2 times a week). Curcumin, ITH12575 and CGP37157 solutions are prepared at 10mM in a DMSO/ethanol mixture (60/40). Then a dilution in a saline solution is carried out to obtain the final desired concentration All treatments were administered intraperitoneally. Tumors were measured twice each week with callipers and tumor volumes were calculated by using the formula [(*π* / 6) × (*w*1 × *w*2 × *w*2)]. Authorisation (Apafis 19933) was given by the regional ethical committee CEEA – 003 (Campus CNRS, Orléans) and MESRI (French Ministry of Higher Education, Research and Innovation).

### Mathematical Models of Tumor Growth

The change in tumor volume over time was described using a generalised logistic model. This model included three parameters: first-order growth rate constant (k_growth_), maximum tumor volume (Vmax) and the power coefficient (γ estimated using nonlinear mixed-effects modelling. This approach aims at quantifying the inter-subject distribution of each model parameter in the population, by estimating their mean and inter-subject standard deviation, as well as quantifying the association of factors of variability (referred to as covariates) and the inter-subject distribution of model parameters. We tested the influence of compounds (CGP37157, ITH12575 and curcumin) as covariates of Vmax because its inter-subject standard deviation was the only one that could be estimated. This influence was tested using the likelihood ratio test (LRT). This analysis was performed using MonolixSuite2019 (Lixoft®, Antony, France).

### Mid-Infrared Spectroscopy

Total blood collected from the xenograft mouse model was analysed at D0 (as the beginning of the experiment) and D40 (as 40 days after treatment began) by MIR spectroscopy by using a Lumos microspectrophotometer (Bruker, Billerica, MA, USA). Acquisitions were made in transmission mode on a ZnSe multi-plate. Spectrum pre-treatments with smoothed second derivatives and vector normalisation were operated. Statistical treatments were performed in the R environment. Statistical models were built with two goals in mind. The first objective was to estimate a decision rule that takes a spectrum as input and returns the probability that the sample belongs to a given group. The second objective was to return a signature of the treatment: a set of wavelengths that characterise the difference between the two compared groups.

The Lasso penalised logistic regression was used because it allows one to deal with both of the aforementioned goals at once; in particular, it is a powerful method for variable selection (49). This method is implemented in the *glmnet* package in the R environment. A k-fold cross validation function was run to select the penalisation constant that controls the sparsity of the estimator.

Successively, the day and the treatment variable were entered as the dependent variable, Y, in the logistic regression model and coded as 1 for the target group and 0 for the others.

To assess predictive performance, the dataset is randomly divided into a training set and a validation set. If the total effect is large enough, Monte Carlo cross-validation is used with 30 repetitions to obtain 30 sets of misclassification errors (50). When the total number of samples was smaller than 20, the leave-one-out approaches performed and leads to one estimated error per observation (51).

We fitted the logistic LASSO regression using the training dataset only and predicted the groups of the test data using the fitted models. Then, classification errors were computed for the test dataset. The cut-off value of the probability for classification, which was needed for calculating the test misclassification errors, was set at 0.5.

The MIR spectral variable assignment (cm^−1^) was as follows: 3320, λ_CH_ stretching in lipids; 3220, λ_OH_ carboxyl groups; 3100–3050, amide B band of proteins; 2813, lipid band tail (aliphatic chains λ_CH2/CH3_); 1750, λ_C=O_ esterified lipids (cholesterol, triglycerides and phospholipids); 1600–1680, amide I band of proteins (sensitive to secondary structure); 1520–1580, amide II band of proteins; 1450, fat deformation λ_CH_; 1380, lipid λ_CH3_ bending; 1350, fat deformation CH_3_; 980, alcohol function CO; and 900, aromatic rings.

### Metabolomic Analysis

Liquid chromatography–high-resolution mass spectrometry (LC-HRMS) was performed after an extraction with 400 µL of methanol from 25 µL of plasma. An Ultimate WPS-3000 UPLC system (Dionex, Germany) coupled to a Q-Exactive mass spectrometer (Thermo Fisher Scientific, Bremen, Germany) and operated in positive and negative electrospray ionisation modes (ESI+ and ESI-, respectively; separate analysis for each ionisation mode) was used for this analysis. Liquid chromatography was performed using a Phenomenex Kinetex 1.7 µm XB – C18 column (150 mm × 2.10 mm) maintained at 55°C. Two mobile phase gradients were used. The gradient was maintained at a flow rate of 0.4 mL/min over a runtime of 20 min. Two different columns were used to increase the metabolic coverage. Accordingly, a hydrophilic interaction liquid chromatography (HILIC) column (150 mm × 2.10 mm, 100 Å) was also used. During the full-scan acquisition, which ranged from 58 to 870 *m*/*z*, the instrument operated at 70,000 resolution (*m*/*z* = 200).

As required for all biological analyses, the pre-analytical and analytical steps of the experiment were validated based on the findings of quality control (QC) samples (mix of all the samples analysed). Coefficients of variation (CV% = [standard deviation / mean] × 100), were calculated from all metabolite data. If a metabolite had a CV > 30% in QC samples, it was excluded from the final dataset. QC samples were analysed at the beginning of the run, every 13 samples and at the end of the run.

The samples were subjected to a targeted analysis, based on a library of standard compounds (Mass Spectroscopy Metabolite Library [MSML®] of standards, IROA Technologies, Ann Arbor, MI, USA). The following criteria were followed to identify the metabolites: (1) retention time of the detected metabolite within ± 20 s of the standard reference; (2) exact measured molecular mass of the metabolite within a range of 10 ppm around the known molecular mass of the reference compound; and (3) correspondence between isotopic ratios of the metabolite and the standard reference (52) (53). The signal value was calculated using the Xcalibur® software (Thermo Fisher Scientific, San Jose, CA, USA) by integrating the chromatographic peak area corresponding to the selected metabolite.

#### Univariate Analysis

The univariate analysis of metabolite levels between groups was realised by using the non-parametric Mann–Whitney test in the XLSTAT software (Addinsoft, Paris, France). *P* < 0.05 was considered significant.

#### Multivariate Analysis

Multivariate analyses were performed as described previously (52)(53) using the SIMCA P+ 13.0 software (Umetrics, Umea, Sweden), which included a principal component analysis (PCA) and an orthogonal partial least squares discriminant analysis (OPLS-DA). The data analysis was preceded by unit variance scaling. The quality of the model was described by the cumulative modelled variation in the X matrix R2 X, the cumulative modelled variation in the Y matrix R2 Y and the cross-validated predictive ability Q2 values. To evaluate the significance of the created model, cross-validation analysis of variance (CV-ANOVA) was applied. The discriminant metabolites named variable importance in projection (VIP) obtained from OPLS-DA were all considered as responsible for the differences between samples.

#### Pathway Analysis

Metabolites of VIP obtained from the final OPLS-DA model were introduced in the pathway analysis module in the MetaboAnalyst platform.

### Quantification and Statistical Analysis

The data are presented as the mean ± standard deviation (SD) or standard error of the mean (SEM) and were analysed using GraphPad 7 (GraphPad Software, La Jolla, CA, USA). In the box plot, the box represents the 25th to 75th interquartile range, the midline in the box represents the median and the solid square box represents the mean. To test single variables between two groups, a paired Student’s t-test or Mann-Whitney test was performed. One-way ANOVA followed by post hoc Tukey’s test was used for multiple comparisons. *P* < 0.05 was considered to be significant.

## Author contributions

Study concept and design: MG, WR and TL Acquisition of data: MG, SI, JB, AR, TP, XZ, JD, DT, RG, HFR, AL, SL, RC, MO, OS, OH, GFH, DTo, TL Analysis and interpretation of data: MG, SI, JB, AR, TP, XZ, DC, JD, DT, VM, HFR, AL, SL, ALP, JFD, PGF, AG, RC, FG, AJGS, PE, OS, OH, GFH, CV, DTo, MT, WR, TL Drafting of the manuscript: MG, SI, JB, PE, OS, CV, DTo, MT, WR, TL Critical revision of the manuscript for important intellectual content: ALL Statistical analysis: MG, SI, JB, VM, PE, DTo, WR, TL Obtained funding: MG, AG, DTo, WR, TL Study supervision: MG, WR, TL

## Acknowledgments

We thank Dr Virginie André and Isabelle Domingo for technical assistance. This project was partly supported by grants on behalf of the following french department committees of Ligue Contre le Cancer Grand-Ouest: 16 (Charente), 37 (Indre-et-Loire), 49 (Maine-et-Loire), 72 (Sarthe) and 85 (Vendée). Maxime Guéguinou is a recipient of a 3-years postdoctoral grant from Région Centre – Val de Loire. Sajida Ibrahim was partly funded by ARFMAD (Association Recherche et Formation dans les Maladies de l’Appareil Digestif). Alison Robert is a recipient of a 3-years doctoral grant from Inserm/ Région Centre – Val de Loire.

Inserm UMR 1069 is leader of Cancéropole Grand-Ouest 3MC network (Marine Molecules, Metabolism and Cancer), member of Région Centre – Val de Loire thematic research consortium RTR MOTIVHEALTH (Molecular and Technological Innovation for Health) and member of CNRS research group APPICOM (Integrative Approach for a multiscale functional analysis of membrane proteins). We acknowledge Inserm, Université de Tours and Région Centre-Val de Loire for their financial supports.

COLOMIN cohort was supported in part by a grant from the association “Sport et Collection” and “Ligue Contre le Cancer, Comités départementaux de la Vienne, Charente et Charente-Maritime” (David Tougeron).

---

**Supplementary Figure 1.**
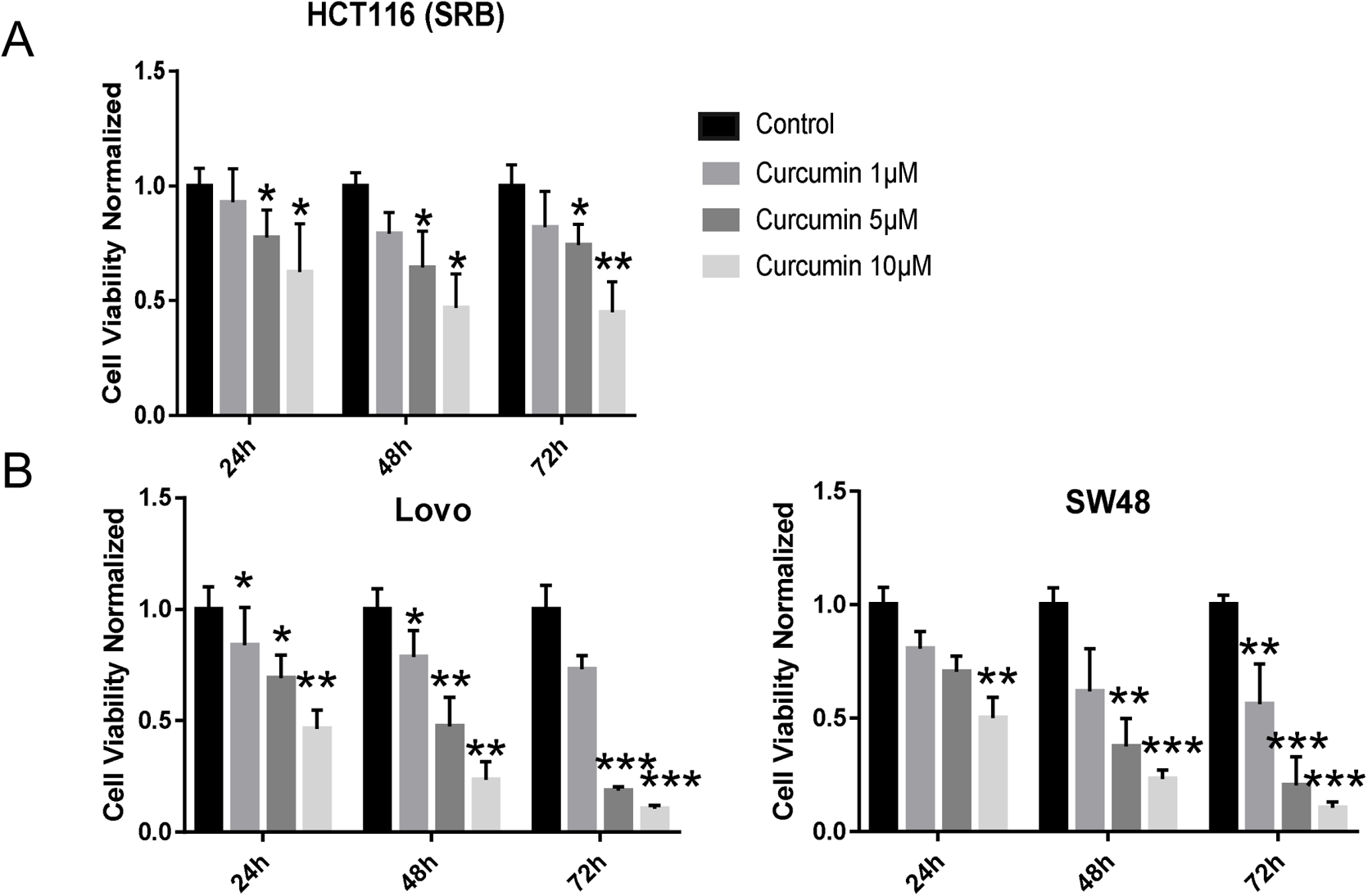

**Supplementary Figure 2.**
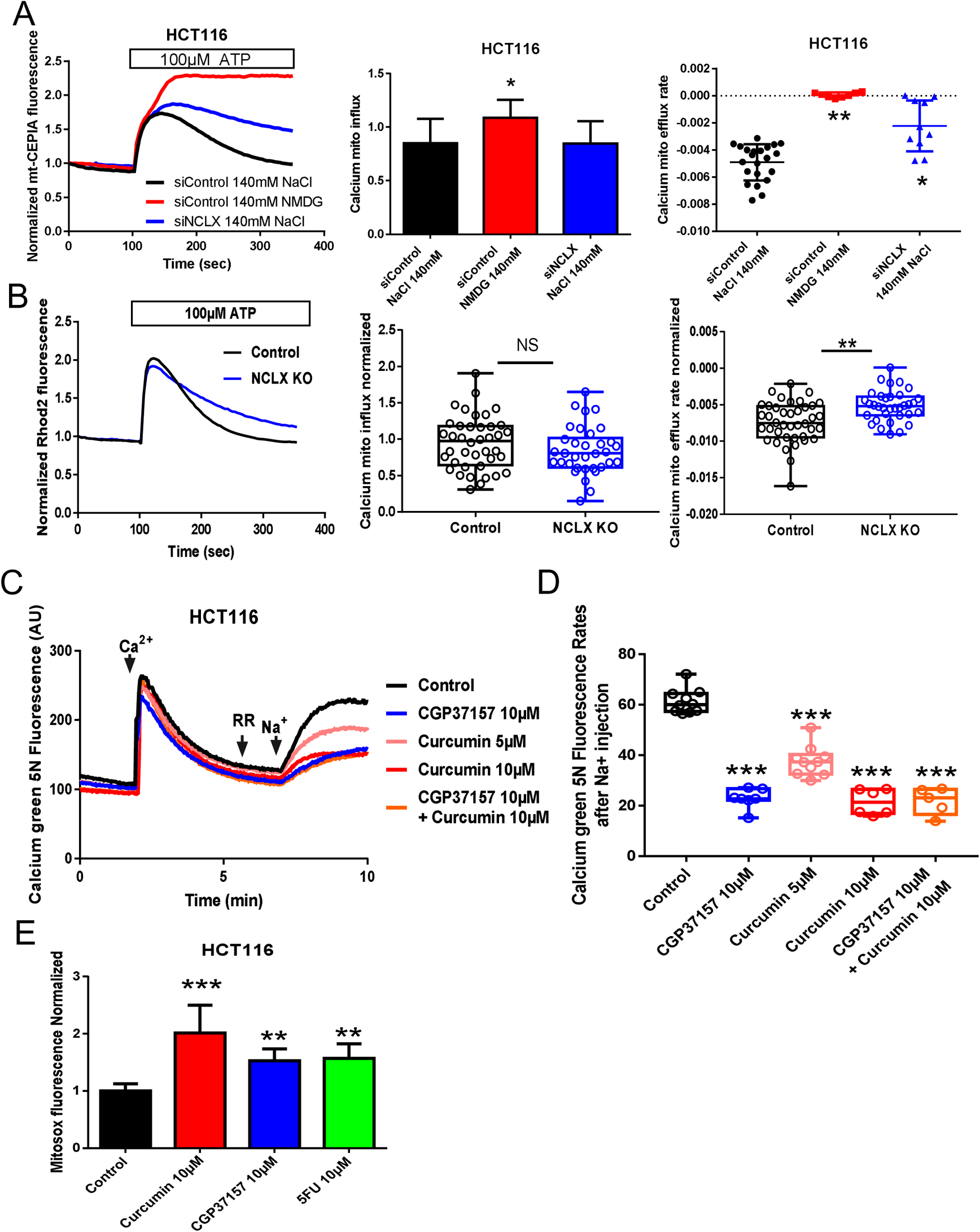

**Supplementary Figure 3.**
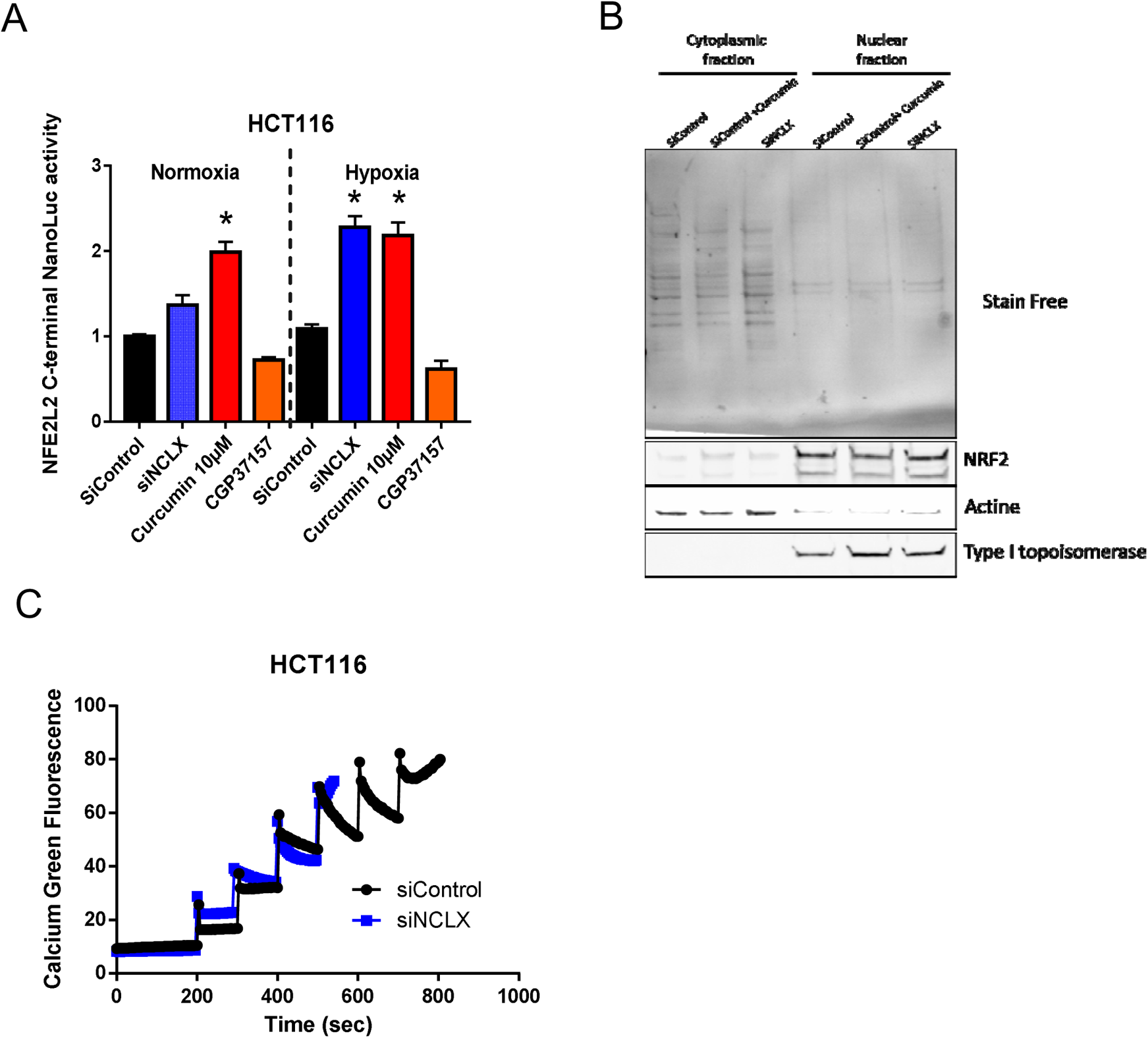

**Supplementary Figure 4.**
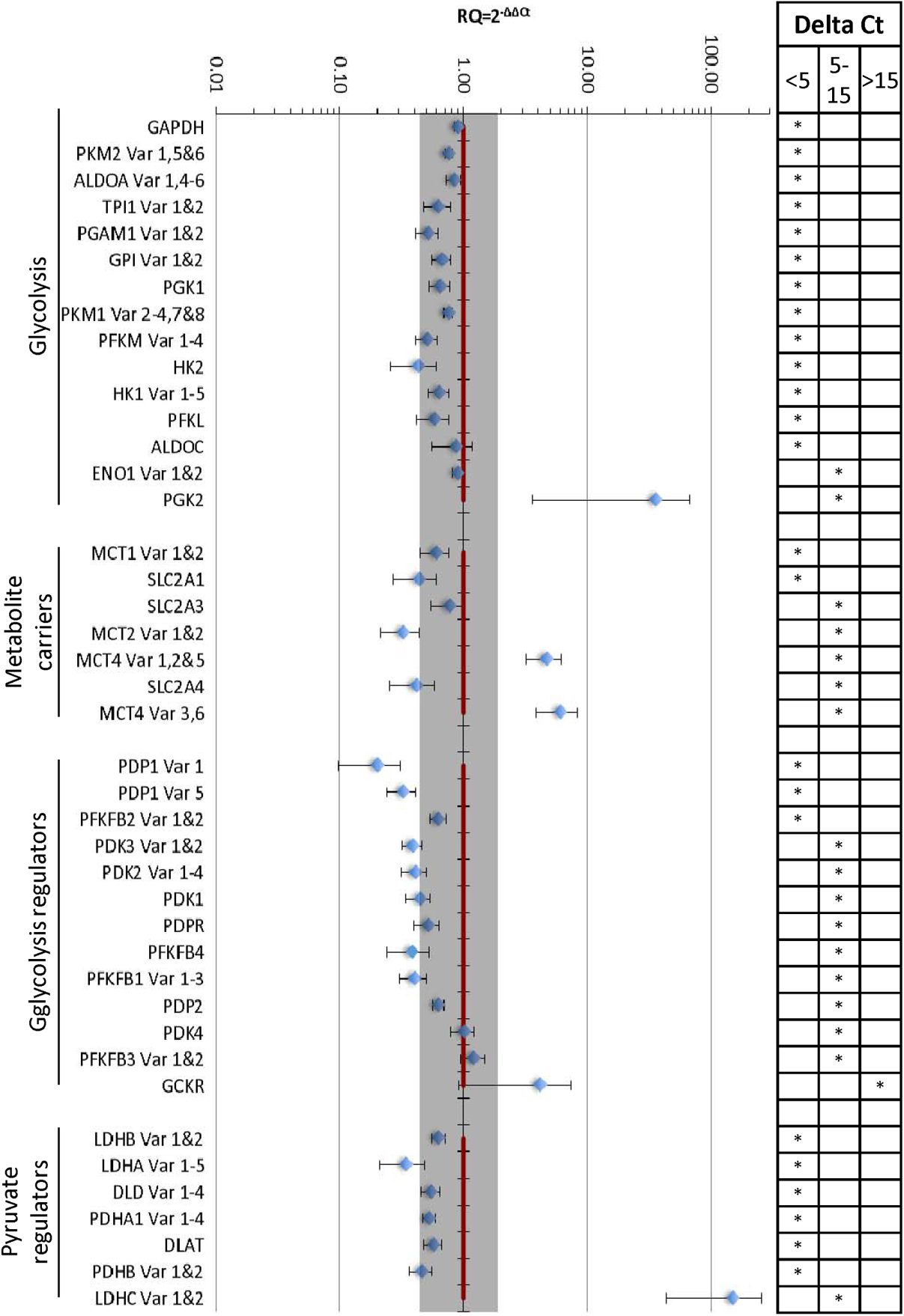
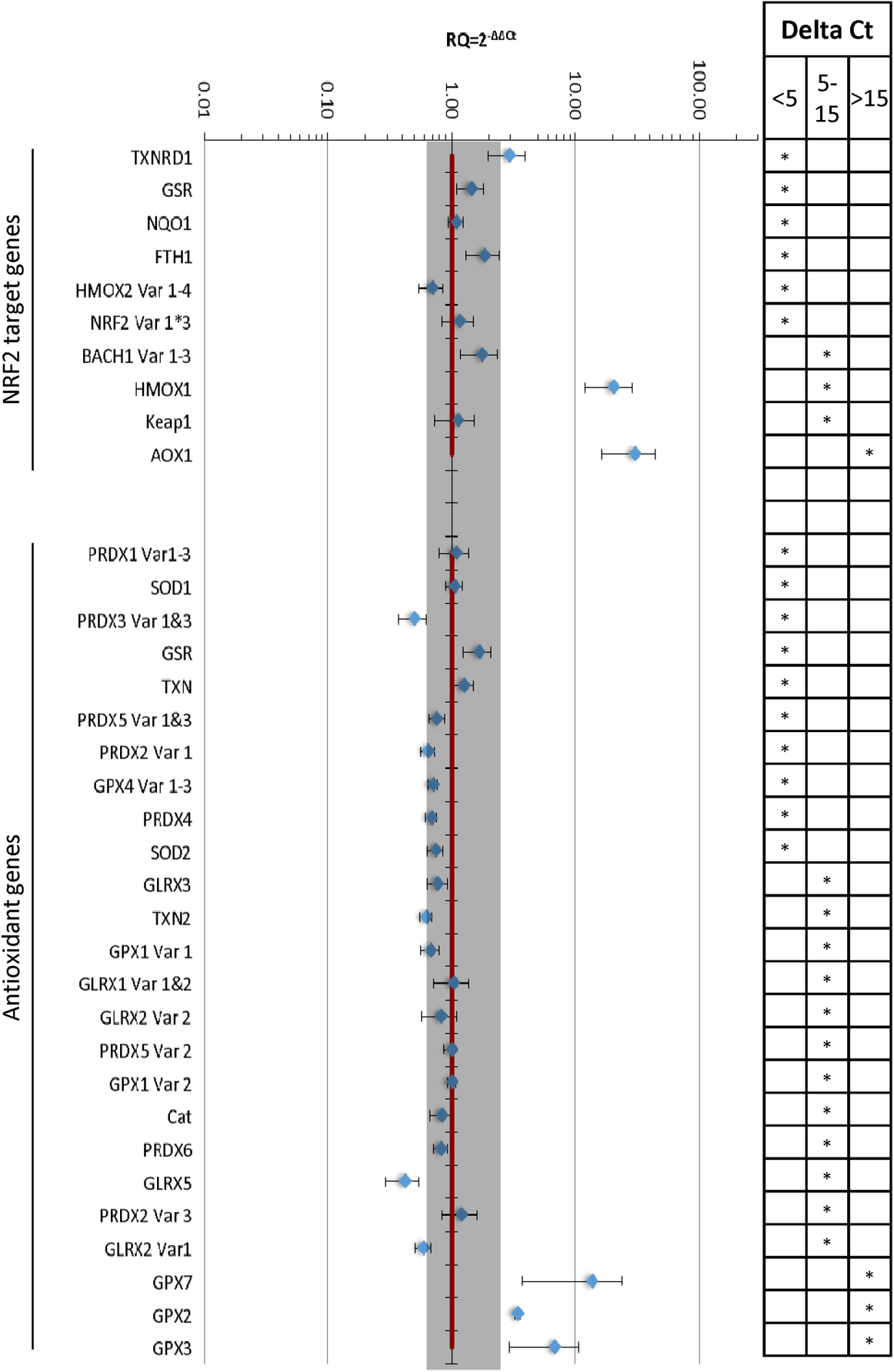

**Supplementary Figure 5.**
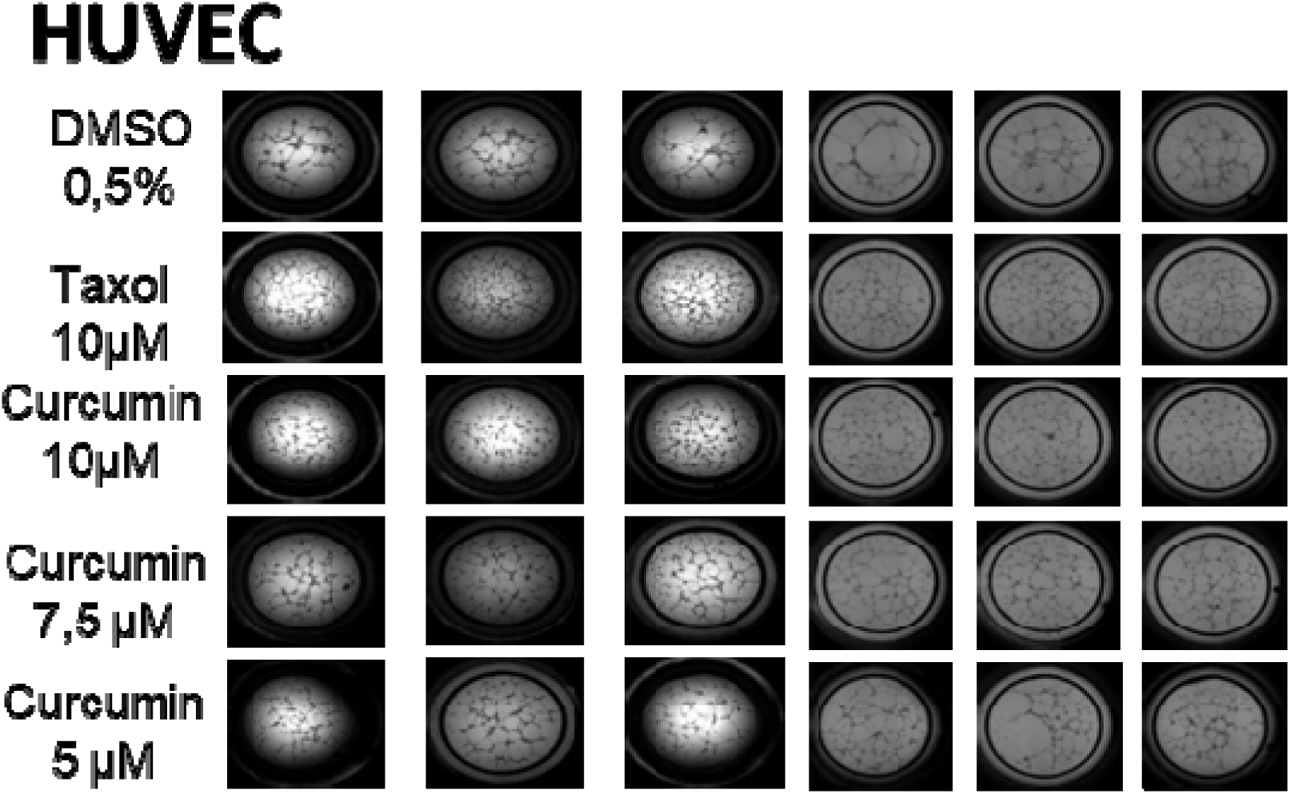

