## Supplementary Tables and Legends for "Curcumin and NCLX Inhibitors Share Anti-Tumoral Mechanisms in Microsatellite-Instability-Driven Colorectal Cancer"

**Supplementary Data**

**Supplementary Table S1: List of primers used for oxidative stress evaluation and genes involved in glycolytic metabolism**

var = splicing variants

| **Gene** | **Forward primer** | **Reverse primer** |
| --- | --- | --- |
| ACTB | attggcaatgagcggttc | cgtggatgccacaggact |
| ALDOA_var 1-4 &6 | tgccagtatgtgaccgagaa | gccttccaggtagatgtggt |
| ALDOC | gaacccgagctgtgcttg | gtacgagtgaggcatggtga |
| AOX1 | ctgagtgttgtaaaccatgcatataa | aatgtccttcacctgggcta |
| BACH1_var1-3 | gcagcagttacttccactcaag | ggttcaaatcctttaactgtcacc |
| CAT | cgcagttcggttctccac | gggtcccgaactgtgtca |
| DLAT | gggtgagaagctaagtgaagga | cagggaccaggatttttgc |
| DLD_var1-4 | cccagaggtagaattccagtca | gagccagcattggaccag |
| ENO1_var 1&2 | tcccaacatcctggagaataa | atgccgatgaccaccttatc |
| FTH1 | gccagaactaccaccaggac | catcatcgcggtcaaagtag |
| GAPDH | agccacatcgctcagacac | gcccaatacgaccaaatcc |
| GCKR | agatgatattcgggctgctc | ggcgatgggtatcacctgt |
| GLRX1 var1&2 | ggcttctggaatttgtcgat | tgcatccgcctatacaatctt |
| GLRX2 var1 | gtggcactcgctggaatc | cgtcgctaaattctccaaagat |
| GLRX2 var2 | gctggtttggagcaggag | ccaaagatgatgatgtattgctct |
| GLRX3 | tcctcaagaaccacgctgt | tgagaagatatcaaaactgctaaactg |
| GLRX5 | gtgataactggggcgttgtt | actcaggcatgcacagca |
| GPI_var1&2 | ccctatgaccagtacctgcac | ttcccattggactccatgtc |
| GPX1var1 | caaccagtttgggcatcag | gttcacctcgcacttctcg |
| GPX1var2 | cccttgtttgtggttagaacg | gagagaagggcagctagaacc |
| GPX2 | gtccttggcttcccttgc | tgttcaggatctcctcattctg |
| GPX3 | cagagatccttcctaccctcaa | ccctttctcaaagagctgga |
| GPX4 var1-3 | tacggacccatggaggag | ccacacacttgtggagctagaa |
| *GPX5_var1&2* | tgctttgtgcaaacaagtcc | ggtgcctttctcgtctttgt |
| *GPX6* | aggcttggcagctcagtatc | gccaacacaatgacaccaaa |
| GPX7 | ccatcctgccttcaagtacc | ttccatctggggctactagg |
| GSR | tgccagcttaggaataaccag | cctgcaccaacaatgacg |
| HK1_var1-5 | cacctgtgaggttggactca | ccaccatctccacgttcttc |
| HK2 | tcccctgccaccagacta | tggacttgaatcccttggtc |
| HMOX1 | gggtgatagaagaggccaaga | agctcctgcaactcctcaaa |
| HMOX2 var 1-4 | ggacttcttgaaaggcaacatta | tgaagtaaagtgccgtggtg |
| KEAP1_var1&2 | accacaacagtgtggagaggt | cgatccttcgtgtcagcat |
| LDHA_var 1-5 | gtccttggggaacatggag | ttcagagagacaccagcaaca |
| LDHB_var1&2 | gatggattttgggggaacat | aacacctgccacattcacac |
| LDHC_var1&2 | gctctgaagactctggacccta | tcagcttgataatttcataggcact |
| MCT1_var 1&2 (SLC16A1) | gtgaccattgtggaatgctg | catgtcattgagccgaccta |
| MCT2 v1-2 (SLC16A7) | gttgacagcgaggcgaat | cttgtttcaggtttcacaggaa |
| MCT4_var 3&6 (SLC16A4) | gggaaggtccaaccttacact | aatagcaaccaggggacca |
| MCT4_var1-2&5 (SLC16A4) | gggaaggtccaaccttacact | ccatcacaaacacattcacca |
| NQO1 | cggctttgaagaagaaaggat | cgcagggtccttcagtttac |
| NRF2 (NFE2L2)_var1-3 | gagacaggtgaatttctcccaat | tttgggaatgtgggcaac |
| PDHA1_var 1-4 | gtccgagaggcaacaaggt | aagtctgcagctccatcagg |
| PDHB_var 1&2 | cggatagaggacacgacca | gtccagtgaaagcgcctct |
| PDK1 | caccaagacctcgtgttgag | acgtgatatgggcaatccat |
| PDK2_var 1-4 | ctggccaacatcatgaaaga | ccaggaggctctggacatac |
| PDK3_var 1&2 | tgtgtgaacagtattacctggtagc | gtttgtctggcgctttgg |
| PDK4 | cagtgcaattggttaaaagctg | ggtcatctgggcttttctca |
| PDP1_var1 | atgttgtcggctccgtgt | tggaacttctgactgggattc |
| PDP1_var5 | tcgggaagaatcgtttggt | aaaaacagttgagttggtgctg |
| PDP2 | ctgagcctgaggtcacatacc | aggccagcacaaggaactta |
| PDPR | gtggcctatcacctctccaa | gccagcacagaacctggtag |
| PFKFB1 var 1-3 | acatggaagccctgcaaat | ggctgagatagttgtgaacatcc |
| PFKFB2_var 1&2 | atctctcggggtgccctat | tgcataggtcatctcttcacaca |
| PFKFB3_var 1&2 | caacagctttgaggagcatgt | gggagcctttcatgttttgt |
| PFKFB4 | ccagatgaagaggacaatcca | tcctcgtaggtcatttcctca |
| PFKL | gcttcgacacccgtgtaact | atgcccatcttgctgctc |
| PFKM_var 1-4 | gccatcagcctttgacaga | ctccaaaagtgccatcactg |
| PGAM1 var1-2 | ggaggggaaacgtgtactgat | agctccatgatagcctcttca |
| PGK1 | cagctgctgggtctgtcat | gctggctcggctttaacc |
| PGK2 | ccagctcctggttcagtca | tcccttcttcctccacatga |
| PKM1_var 2-4 & 7-8 | cagccaaaggggactatcct | cctcagcctcacgagctatc |
| PKM2_var 1&5-6 | cagccaaaggggactatcct | caaataattgcaagtggtagatgg |
| PRDX1 var1-3 | cactgacaaacatggggaagt | tttgctcttttggacatcagg |
| PRDX2 var1 | gccttccagtacacagacgag | gttgggcttaatcgtgtcact |
| PRDX2 var3 | gcaactcagatgcaactctatctact | tgaactggagtttccatcttcat |
| PRDX3 var1&2 | ctggacaccggattctccta | gggtgatctactgatttaccttctg |
| PRDX4 | gcacctaagcaaagcgaaga | aaattctccatcgatcacagc |
| PRDX5 var1/3 | tcctggctgatcccactg | atgccatcctgtaccaccat |
| PRDX5 var2 | cacccctggatgttccaa | ggacaccagcgaatcatctagt |
| PRDX6 | caatagacagtgttgaggaccatc | tttctgtgggctcttcacaa |
| RPL13A | caagcggatgaacaccaac | tgtggggcagcatacctc |
| SLC2A1 | ggttgtgccatactcatgacc | cagataggacatccagggtagc |
| SLC2A3 | gccctgaaagtcccagattt | ttcatctcctggatgtcttgg |
| SLC2A4 | ctgtgccatcctgatgactg | cgtagctcatggctggaact |
| SOD1 | gcatcatcaatttcgagcag | caggccttcagtcagtcctt |
| SOD2 | tccactgcaaggaacaacag | taagcgtgctcccacacat |
| SOD3 | ctctcttttcaggagagaaagctc | aacacagtagcgccagcat |
| SULF1 | ccaatgcttcccaacacata | gcagcattggtcctgtgtact |
| TPI1_var1&2 | gttgggggaaactggaagat | tagggggagcacaaaccac |
| TXN | ttacagccgctcgtcaga | ggcttcctgaaaagcagtctt |
| TXN2 | gagacaccagtggttgtgga | gcttggccaccatcttctc |
| TXNRD1 | tcaccccagttgcaatcc | ggttggaacattttcatagtcaca |

**Supplementary Figure 1: Cell Viability of MSI-Colorectal Cancer Cells**

A/ HCT116 cells were incubated with various concentrations of curcumin (0–10 µM) for 24, 48 or 72 h and cell viability was measured by using SRB assay. The results are expressed as the mean ± standard deviation of 3–5 independent experiments (ANOVA followed by Tukey’s post hoc multiple comparisons; **P* < 0.05, ***P* < 0.01).

B/ Lovo or SW48 cells were incubated with various concentrations of curcumin (0–10 µM) for 24, 48 or 72 h and cell viability was measured using the MTT or SRB assay. The results are expressed as the mean ± standard deviation of 3–5 independent experiments (ANOVA followed by Tukey’s post hoc multiple comparisons, **P* < 0.05, ***P* < 0.01 and ****P* < 0.001).

**Supplementary Figure 2: Characterization of NCLX on Calcium or ROS Signalling**

A/ mtCa^2+^ responses following application of ATP (100 μM, as in Figure 3A) was measured in HCT116 cells (siControl or siNCLX with 140 mM NMDG). The mtCa^2+^ influx (peak) (left) and the rates of mtCa^2+^ efflux are shown (right) are shown as the mean of three independent experiments with 8–22 measurements (Mann–Whitney test, ****P* < 0.01).

B/ mtCa^2+^ responses following application of ATP (100 μM, as in Figure 3A) were measured in HCT116 control (NCLX +/+) or NCLX KO (-/-) cells. The average mtCa^2+^ influx (peak) (left) and rates of mtCa^2+^ efflux (right) are shown (n = 9, Mann-Whitney test, ***P* < 0.01)

C/ Average trace of permeabilized HCT116 treated with vehicle, CGP37157 or Curcumin at T0 min cells in intracellular Na+-free buffer ( 130 mM KCl, 10 mM Tris-MOPS (pH 7.4), 10 μM EGTA-Tris, 1 mM KPi, 5 mM malate, 5 mM glutamic acid, 1 μM Ca2+ green 5N). Additions were made as indicated by arrows: Ca2+ (10 μM; 2min), ruthenium red (RR) (0.5 μM; 6min), NaCl (10 mM; 7min)

D/ Quantification of Ca2+ efflux rate as shown between 7min and 8.5 min. Data shown are Box-plot efflux rate ± SD (N=5-10). ***p < 0.001, one way ANOVA with Dunnett's multiple comparisons test.

E/ The effect of CGP37157, curcumin or 5-fluorouracil (5-FU) pre-treatment on mtROS levels in HCT116 cells were recorded. The mtROS level was measured using the MitoSOX red dye.

**Supplementary Figure 3: Characterization of NCLX on NRF2 expression/translocation and PTP activation**

A/ HCT116 NFE2L2-C-terminal Luc cells were incubated under normoxic or hypoxic conditions for 48 h. Four hours before the end of incubation, the cells were treated with CGP37157 or curcumin. After incubation, luciferase expression was determined. The values represent the mean ± standard deviation of 3–6 independent experiments with 11–61 measurements.

B/ Representative western blot showing the effect of curcumin treatment or siNCLX on NRF2 nuclear translocation.

C/ Extramitochondrial calcium traces with isolated mitochondria from HCT116 siControl and siNCLX using the calcium indicator dye Calcium Green-5N with different. calcium addition.

**Supplementary Figure 4 (a and b): Gene Expression Profiling of Metabolic Pathways**

Relative quantification (RQ) gene expression of glycolysis, metabolite carriers, glycolysis regulators, pyruvate regulators, NRF2 target, antioxidant genes was evaluated by the 2^-ΔΔCt^ method on HCT116 treated or not with curcumin (n = 4 independent experiments).

**Supplementary Figure 5: Tube Formation Assay**

HUVEC (human umbilical vein endothelial cells) tube formation assay with 10 µM Taxol and 5–7.5 and 10 µM of curcumin. Treatment with curcumin completely abrogated the network, the branches and the bridges between cells. The images are representative examples.
